# Combined mRNA and protein single cell analysis in a dynamic cellular system using SPARC

**DOI:** 10.1101/749473

**Authors:** Johan Reimegård, Marcus Danielsson, Marcel Tarbier, Jens Schuster, Sathishkumar Baskaran, Styliani Panagiotou, Niklas Dahl, Marc R. Friedländer, Caroline J. Gallant

## Abstract

Combined measurements of mRNA and protein expression in single cells enables in-depth analysis of cellular states. We present single-cell protein and RNA co-profiling (SPARC), an approach to simultaneously measure global mRNA and large sets of intracellular protein in individual cells. Using SPARC, we show that mRNA expression fails to accurately reflect protein abundance at the time of measurement in human embryonic stem cells, although the direction of changes of mRNA and protein expression are in agreement during cellular differentiation. Moreover, protein levels of transcription factors better predict their downstream effects than do the corresponding transcripts. We further show that changes of the balance between protein and mRNA expression levels can be applied to infer expression kinetic trajectories, revealing future states of individual cells. Finally, we highlight that mRNA expression may be more varied among cells than levels of the corresponding proteins. Overall, our results demonstrate that mRNA and protein measurements in single cells provide different and complementary information regarding cell states. Accordingly, SPARC can offer valuable insights in gene expression programs of single cells.

## INTRODUCTION

Advances in single cell analysis are impacting the scale and resolution at which we investigate biological systems. The majority of these advances are focused on the application of single cell RNA sequencing (scRNAseq). However, transcriptomes may not fully reflect cell states and cellular signal transmission. scRNAseq provides comprehensive snapshots of gene expression, but gene transcription is stochastic and characterized by transcriptional bursts of varying rates and sizes^1^, and half-lives of mRNA molecules vary significantly between genes^2^. Sampling from low number of molecules, variable efficiency to convert mRNA to cDNA and PCR amplification bias during sequencing library preparation also contribute to noisy expression data^3^.

In contrast to mRNA, proteins are more stable, and typically present in orders of magnitude higher amounts within the cell^2^, reducing chance fluctuations of their levels. Moreover, proteins have more direct roles in maintaining cellular functions compared to transcripts. While bulk sample measurements generally report good correlation between mRNA and protein expression^2,4-6^, there has been little insight into the extent of the agreement at cellular resolution in systems at steady-state or undergoing a dynamic transition. Therefore, we argue that combined mRNA and protein single cell measurement approaches are necessary to better understand cellular states, the causes and consequences of cell-to-cell variability, and to decipher regulatory circuits and pathways.

A number of approaches are emerging to simultaneously measure both mRNA and protein at single cell resolution. These include measurement of cell surface proteins and mRNA in droplet microfluidic-based analysis platforms^7,8^, targeted mRNA and protein detection^9-12^, and mRNA and targeted protein measurement for a limited number of cellular proteins (n=6) in fixed cells^13^. We describe an approach, herein called single-cell protein and RNA co-profiling (SPARC), which enables measurement of global mRNA and high multiplex, targeted cellular proteins in single cells **(Figure 1A).** mRNA levels were recorded using a modified Smart-seq2 protocol^14^ enabling sensitive expression measurements of the full-length transcripts. Proteins were measured using multiplex, homogeneous protein extension assays (PEA)^12, 15^, an affinity-based protein detection method that allows scalable protein detection in fresh cell or tissue lysates without a need for prior fixation.

**Figure 1:**
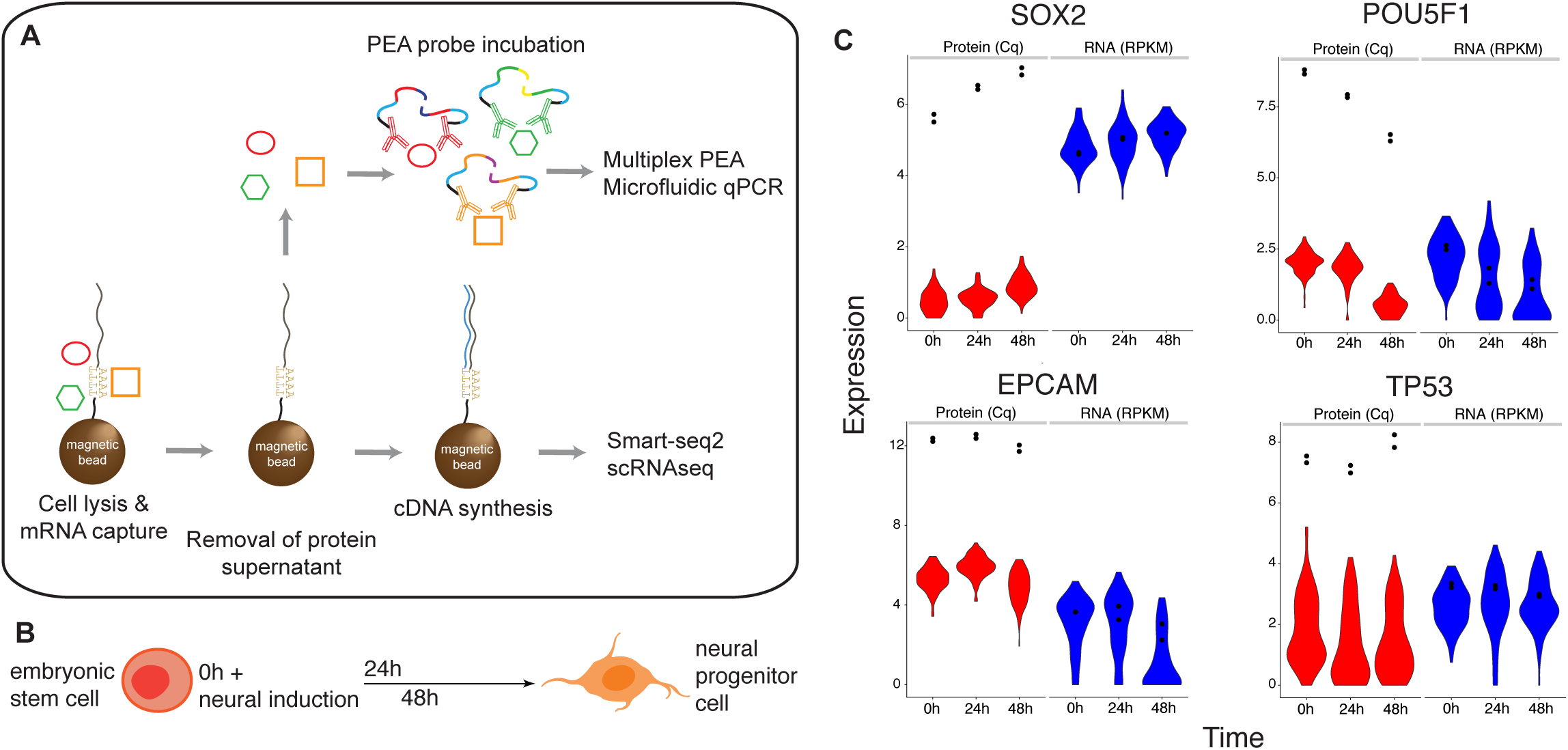
SPARC method overview. The SPARC procedure. Single cells are isolated and lysed in the presence of oligo-dT conjugated magnetic beads. Following oligo-dT mRNA hybridization, the protein-containing supernatant is removed for subsequent multiplex proximity extension analysis (PEA) and the mRNA is processed using a modified Smart-seq2 approach. **(B)** Overview of the cellular model analyzed using SPARC. Human embryonic stem cells were analyzed in culture (0h) and following directed neural induction (24h and 48h). **(C)** Example mRNA (blue) and protein (red) expression in single cells (violin plots) or replicate 100 cell population control (black dots) measured at 0h, 24h and 48h post-neural induction.

PEA is a member of a class of proximity-based assays that require two binding events in order to generate a DNA reporter molecule. In turn, the reporter can be detected and quantified using various DNA detection technologies, including quantitative PCR and sequencing. Here, proteins are detected using pairs of antibodies conjugated with oligonucleotides whose free 3’ ends are pairwise complementary. When cognate antibody pairs bind their target protein, the attached oligonucleotides are brought in proximity and can be extended by polymerization to create amplifiable DNA reporter molecules. A multiplex readout is achieved by decoding extension-generated DNA reporters by real-time PCR using primer pairs specific for cognate pairs of antibody conjugates. The requirement for pairwise protein detection ensures assays equivalent in quality to sandwich immunoassays, while allowing for simultaneous measurement of many proteins in each reaction. Moreover, PEA assays do not require fixation of the cell, capture of target proteins on solid supports or wash steps following the addition of the oligo-conjugated antibody probes to the cell lysate.

Here, we used SPARC to investigate to what extent the amounts of a transcript are predictive of the levels of the corresponding protein in cells at steady-state or undergoing a state-transition. Specifically, we measure mRNA and protein in human embryonic stem cells (hESCs) unperturbed or at fixed time points after induction of directed neuronal differentiation **(Figure 1B)**. We also investigate the effects of biological inferences by analyzing scRNAseq data alone or integrated with the targeted protein expression data and measure the agreement between mRNA and protein expression variation.

We show that mRNA expression fails to accurately reflect protein abundance at the time of measurement in cells, either at steady-state or undergoing neuronal differentiation, although the direction of mRNA and protein expression changes are primarily in agreement during differentiation. We further show that changes of the balance between protein and mRNA expression levels can be applied to infer the kinetic trajectories, or so-called velocity vectors, of single cells^16^, revealing the future state of individual cells. Moreover, we demonstrate that measurements of transcription factors at the protein level better predict their downstream effects than measurement of the corresponding transcripts. Finally, we provide evidence that gene expression variation is not always in agreement for mRNA and protein.

With SPARC, we present a powerful approach to quantitatively measure mRNA and protein in single cells and show how protein measurements greatly aid the analysis of gene expression variation, cell states and cellular regulatory mechanisms.

## RESULTS

### Single cell mRNA expression data

The Smart-seq2 scRNAseq method adapted for the SPARC protocol includes a number of steps that differ from the published protocol (Picelli et al., 2014). The key differences include additional detergents in the cell lysis buffer in order to ensure access to nuclear proteins, the use of oligo-dT conjugated T1 Dynabeads in order to immobilize the poly(A) mRNA fraction, and exclusion of the 72°C heat step before the reverse transcription reaction to avoid denaturing cellular proteins (Methods). We compared the expression data prepared with the reference Smart-seq2 method (0h, n = 67 cells) and SPARC (0h n = 85 cells). The cells were sorted and processed for analysis in parallel and the sequencing was done in the same lane of the sequencing flow cell. Only SPARC data was collected for cells during early differentiation (24h n = 76 cells, and 48h n = 86 cells).

Overall, the scRNA-seq data using SPARC was of high quality and highly comparable to Smart-seq2 (Pearson correlation coefficient = 0.90) **(Figure S1)**. Some minor differences were observed for numbers of genes (n = 3308 for SPARC vs. 3746 for Smart-seq2) or pseudogenes (n = 113 vs. 62), the fraction of reads in introns (n = 0.43 vs. 0.13) and average length of detected genes **(Figure S1)**. We attribute the greater intron capture with SPARC to lysis conditions providing greater access to nuclear content and the omission of a heating step before the oligo-dT primed reverse transcription. Specifically, we may preferentially access and capture unspliced, nascent mRNA before they mature and are fully coated with RNA binding proteins^17^. Nonetheless, the QC analysis showed that the SPARC mRNA data was of high quality and highly reproducible, allowing us to proceed with the combined mRNA and protein analysis.

### Single cell protein expression data

We developed an exploratory multiplex PEA panel for single cell analysis in collaboration with Olink Proteomics, involving 96 proteins and focused on cellular proteins of interest for our investigation. The panel includes proteins across different functional groups related to, for example, pluripotency, neurogenesis, cell cycle phase and metabolic functions **(Table S1).** The PEA protein panel was developed for application across different cellular models, and therefore not all proteins were expected to be detectable at the single cell level in hESCs. Of the 92 proteins in the panel, 87 proteins had detectable levels in the 100 cell control, and 89 proteins in single cells **(Figure 1C) (Table S1) (Figure S2).**

We proceeded to investigate the major sources of variation using PCA for the mRNA and protein expression data sets. Using RNA expression data, the three factors most strongly contributing to variation between the sampled cells were the numbers of detected genes, developmental state and cell cycle phase. For the protein data, the most important factors were total protein abundance (Methods) and developmental state. We also observed a strong positive correlation between numbers of mRNAs or proteins detected and the cumulative sum of protein expression for individual cells, and a weak positive correlation between numbers of genes detected at the level of mRNA and protein. We hypothesize that these correlations reflect the relationship between the amount of RNA or protein and cell size, with larger cells containing more absolute numbers of molecules^18^ **(Figure S3A**,**B)**. Indeed, we observed higher protein levels as measured by protein sum in G2 versus S or G1 cells for cells FACS sorted by cell cycle phase **(Figure S3C).** In order to minimize the effects of cell cycle phase, cell size and mRNA capture efficiency on the mRNA expression variation, the data was normalized for variation related to the cell cycle and the number of genes detected. For the protein measurements, the data was normalized according to cumulative protein sums.

To reduce the dimensionality of the data and align the cells along a trajectory based on similarities in their expression patterns, we next performed tSNE and pseudotime analysis. We applied SCORPIUS^19^, a single trajectory inference method, to order the dynamic cells along a progression from undifferentiated to a more differentiated state. The analysis was done separately on normalized mRNA and protein expression data. The cells were largely ordered and grouped according to the sampled time points (0h, 24h and 48h) (**Figure 2A,B)** and the cell order was very similar whether the trajectory was determined based on mRNA or protein expression data (Pearson correlation coefficient r = 0.82) **(Figure 2C).** The results highlight that both the mRNA and protein data generated with SPARC recapitulate the expected dynamic chanages of the model cell system.

**Figure 2:**
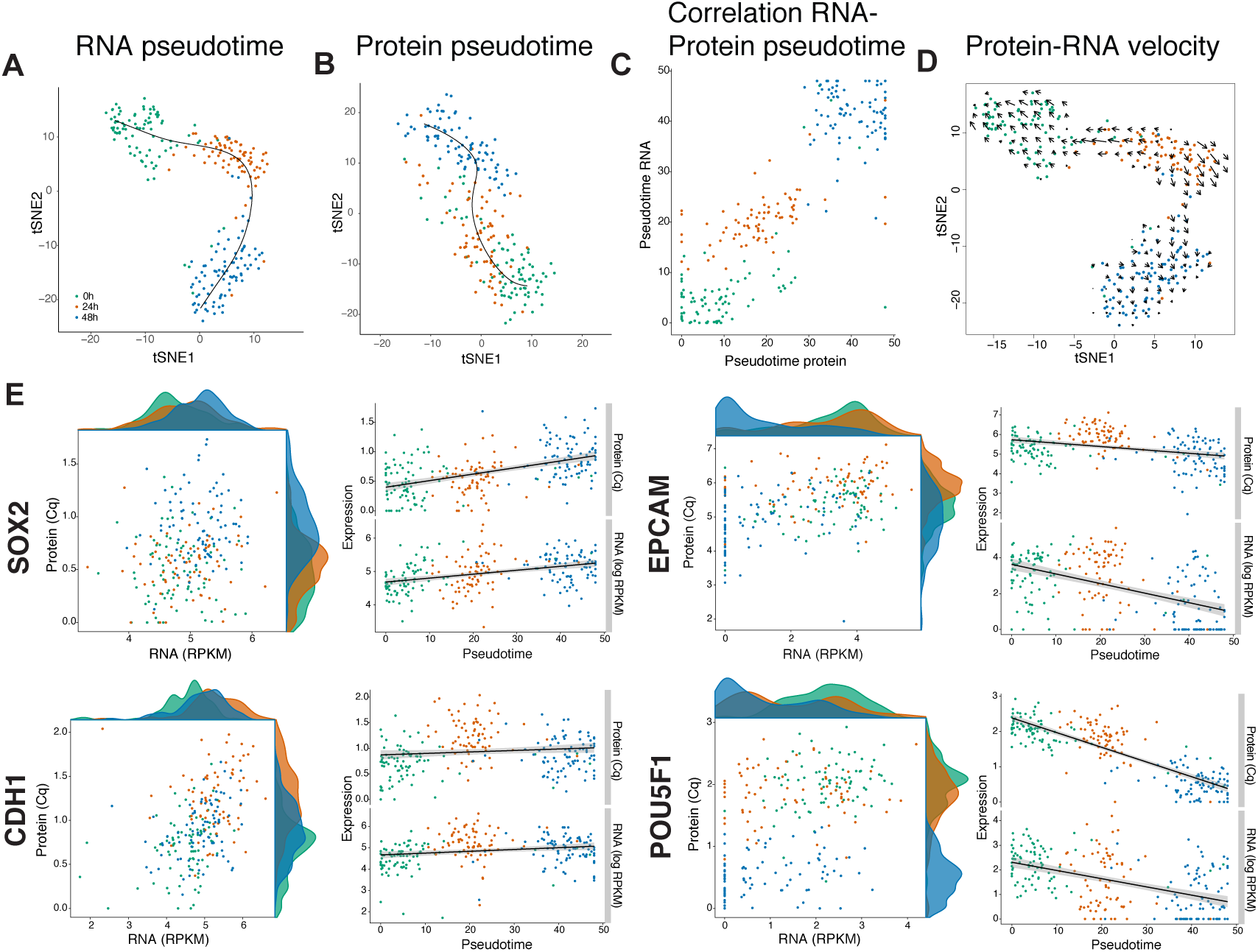
Combined mRNA and protein expression. tSNE plot and pseudotime analysis of the **(A)** normalized mRNA and **(B)** protein expression data. **(C)** Comparison of the cell pseudotime-ordering if calculated based on either mRNA or protein expression data. **(D)** Protein-RNA velocity vectors plotted on the tSNE plot as determined by the normalized mRNA expression. **(E)** Example mRNA and protein expression data plotted for SOX2, EPCAM, CDH1 and POU5F1. The data is shown either as RNA expression vs. protein expression scatter plot with respective density plots included on each axis, or as cells ordered along RNA pseudotime with either RNA and protein expression plotted on the y axis. Both the RNA and protein data are plotted as log2. Color corresponds to time point in all plots 0h (green), 24h (orange) and 48h (blue).

### Relation of RNA and protein data in single cells at steady-state

We next investigated the co-expression of mRNA and protein in cells measured at the 0h time point, purportedly at steady-state. We define cells as being in steady-state when the mean protein or mRNA levels remain relatively constant over several hours ^6,5^. For investigation of correlation, we focused on within gene correlations i.e. the variation of a gene’s mRNA and protein concentrations across single cells^20^. We focused on genes where we detected the protein at a level of > 3 Cq over background in the 100 cell population control in at least one time point.

In agreement with previous studies, we found that levels of mRNA expression are generally a poor predictor of protein expression in single cells (Pearson correlation coefficient between −0.12 to 0.40), and that the scaling and the extent of the relationship between mRNA and protein is gene-specific^9^ **(Figure 2E and Figure S4)**. We measured a limited set of genes across many different functional classes (e.g. transcription factors, metabolic genes), and therefore we do not attempt to make generalized conclusions about mRNA-protein relationships at the gene function level.

For genes where we detect both the mRNA and protein in the majority of the cells, the molecules exist in very different dynamic expression ranges – with the mRNA consistently spanning a much broader range of expression levels compared to protein (e.g. EPCAM, CASP3) **(Figure S4**). We also observed that protein level measurements appear to provide a more stable representation of a gene’s expression state with protein expression detected in the majority of cells measured, whereas mRNA expression detection was more variable, with a fraction of cells showing no detectable expression. This could be because the gene was not expressed at detectable levels at the time of measurement, or the mRNA may have failed to be reliably captured and measured, a common technical problem in scRNAseq protocols **(Figure S4)(Supplemental Results)**.

For some of the genes we measured, the PEA assay did not to detect the protein despite high RNA expression (e.g. PARP1) as the assays did not have single cell sensitivity in the cells investigated, or that the level of detection of the PEA probe is at the limit of detection and results are therefore only qualitative **(Table S2).** Examples of the latter include the detection of the cell cycle markers CCNA2 and AURKA where we expect peak expression during the cell cycle phase G2. Indeed, this is what what we observe, both in single cells predicted to be in G2 (data not shown) and in a separate experiment where cells were sorted by cell cycle phase **(Figure S3C)**. Accordingly, some of the proteins that we could could not detect are likely not or very lowly expressed at specific cell states.

### Relation of RNA and protein levels in single cells during a dynamic change

Next, we investigated the agreement of mRNA and protein expression during early time-points of directed neuronal differentiation **(Figure 1B)**. As with the cells at steady-state, mRNA expression is generally a poor predictor of protein expression. Despite the poor relationship, we were interested to see whether the directional changes of mRNA and protein abundances are in agreement in cells ordered along pseudotime as defined by the mRNA expression data (pseudotime_RNA_) **(Figure 2)(Figure S5).** To test whether the expression levels for both mRNA and protein level show the same directional changes, we applied a linear model for expression over time for both the RNA log-transformed RPKM values and for protein Cq values. The resulting linear model had a low but significant Pearson’s r-square value = 0.20 (p-value <0.0005) **(Figure S6)**. Therefore, the directional changes of mRNA and protein abundances are in general agreement when measured over differentiation time.

In our data, we observed clear examples of temporal delay of gene expression at the level of mRNA and protein expression. This is well demonstrated by POU5F1, a transcription factor that is rapidly turned off in hESCs upon directed neuroectodermal differentiation^21^. When we order the single cells along pseudotime_RNA_ and plot POU5F1 gene expression, we observed that both POU5F1 mRNA and protein show concordant downward trends in expression **(Figure 2E).** However, we note that the protein, but not the mRNA, is detectable in many cells assigned to late pseudotime_RNA_ (corresponding primarily to the cells measured at 48h). We attribute the time-dependent increase in mRNA-protein expression variability as an effect of gene down-regulation and the differential stabilities of mRNA versus protein^22^. Long protein half-lives will enable proteins to be present in a cell long after repression of gene transcription, especially so if targeted protein degradation is not accelerated. We made similar observations for other genes that are down-regulated upon differentiation, including EPCAM. Accordingly, we motivate that when analysing expression data from a cellular system undergoing dynamic changes, the lag-times in mRNA and protein expression should be accounted for in terms of assessing the functional state of the cell at the time of measurement.

Overall, the results highlight that while mRNA expression abundances are not predictive of protein abundances at the time of measurement, the differences can be reconciled when we resolve mRNA and protein expression over a (pseudo)temporal scale, and therefore take into account the temporal elements of gene regulation, including the lag times between transcription and translation, and the different half-lives of mRNA and protein molecules. Accordingly, these results also suggest that mRNA and protein measurements in single cells are not redundant but provide different information regarding the cell state at the time of measurement.

### RNA and protein velocity

In RNA sequencing data, the balance of unspliced and spliced mRNA abundance have been used as an indicator of the future state of mature mRNA abundance, and therefore the future state of the cell ^16, 23^. We hypothesize that in a similar fashion, future cell states as indicated by proteome profiles can, to a certain degree, be predicted from current transcriptome states. In order to explore this assumption, we applied the analysis program for RNA velocity as described by La Manno et al^16^ separately to the mRNA and to the combined mRNA and protein expression data. Our scRNAseq protocol targets polyadenylated, mature spliced mRNA via enrichment with oligo-dT primers. However, there is a consistent fraction of reads mapping to introns, and these reads are attributed to secondary priming within the primary transcripts^16^. In agreement with previous work, the amounts of intron and exon reads are correlated, indicating that introns represent unspliced, precursor mRNA (data not shown).

We first explored whether analysis of our mRNA expression data results in so called RNA velocity vectors – the first time derivative of the spliced mRNA abundance – that are consistent with the direction of differentiation in our cell model. The vector prediction range is on the order of hours^16^, whereas we are sampling every 24h. Therefore, there was a risk that our long sampling intervals would make it difficult to capture meaningful expression dynamics in our data. However, the resulting RNA velocity vectors recapitulate the sampling time and the direction of the differentiation in our cell model **(Figure S7A).**

Next, we asked whether we could perform the velocity analysis by assigning the mRNA expression data as the “unspliced”, immature mRNA input and protein abundance as the “spliced”, mature mRNA input. With this analysis, we aimed to further confirm that the protein expression data is consistent with the biological model, and to further evaluate the relative use of mRNA and protein expression measurements to predict the future states of the cells. We show that the use of combined protein-mRNA expression results in a velocity map **(Figure 2D)** that is consistent with the RNA velocity results **(Figure S7A)**. To verify that RNA-protein velocity results are not driven by changes of total protein abundance, we performed a permuted control and coloured the cells according to their total protein abundance (data not shown). We also permuted the intron count per gene between samples to test if the embedding, i.e. the sample location in the tSNE plot generated in Seurat^24^, could influence the direction of the arrows **(Figure S7B)**. Neither analysis gave an indication that total protein abundance or the pre-defined location of the samples are the main drivers of the velocity vectors. Together, the results show that mRNA abundance can be used to predict future protein abundance in a dynamic system, and importantly, they also highlight the limitations of mRNA expression data to predict protein abundances in the cells at the time of measurement.

### Protein vs. mRNA expression levels correlate better with their trans-regulatory targets

To date, a major limitation of regulatory network inference analysis is the requirement for very high numbers of replicate observations^25^. Single cell analysis experiments produce hundreds or thousands of independent measurements and provide an opportunity to use single cell data to power the analysis of causal gene regulatory network analysis. Despite the increase in power, the use of scRNAseq remains a challenge, in part due to the stochastic nature of RNA transcription and the technical limitations of the sequencing protocols.

We were interested to determine whether integrating protein-level expression data with the mRNA expression would help decipher gene regulatory networks. Our reasons to expect an added value of protein measurements include the observation that single cell mRNA and protein sets are concordant but not redundant, the circumstance that protein expression measurements show lower biological and/or technical-associated cell-to-cell variation than mRNA measurements, and the expectation that protein expression levels of gene regulatory effectors such as transcription factors should be a closer representation of their functional activity at time of measurement.

To specifically address the question of how protein measurement can complement transcriptomics in analyses of gene regulatory networks, we asked whether expression changes at the level of RNA or protein of a transcription factor better predict downstream changes to its targets. We decided to focus our analysis here on POU5F1 as it is essential stem cell factor, it positively and negatively regulates many genes, and that it is rapidly turned off upon differentiation. We expect to observe both (a) POU5F1-POU5F1_target_ co-variation in steady-state cells (0h) as the cells manage their pluripotent state, and (b) POU5F1-POU5F1_target_ correlation as POU5F1 is turned off upon differentiation (0h, 24h, 48h). We first investigated the relationship between either POU5F1_RNA_ or POU5F1_protein_ expression levels and the mRNA expression levels of a subset (n = 7) of well-established downstream POU5F1 trans-regulatory targets in ESCs (Methods) **(Figure 3A, B)**. We did the analysis focusing either on the cells at steady-state (0h) or including all time points analyzed (0h, 24h, 48h). Overall, the correlation of mRNA expression to POU5F1 target genes is stronger for POU5F1 protein values than for POU5F1 mRNA expression. This is true for analysis of cells either at steady-state and cells undergoing differentiation **(Figure 3A, B).**

**Figure 3:**
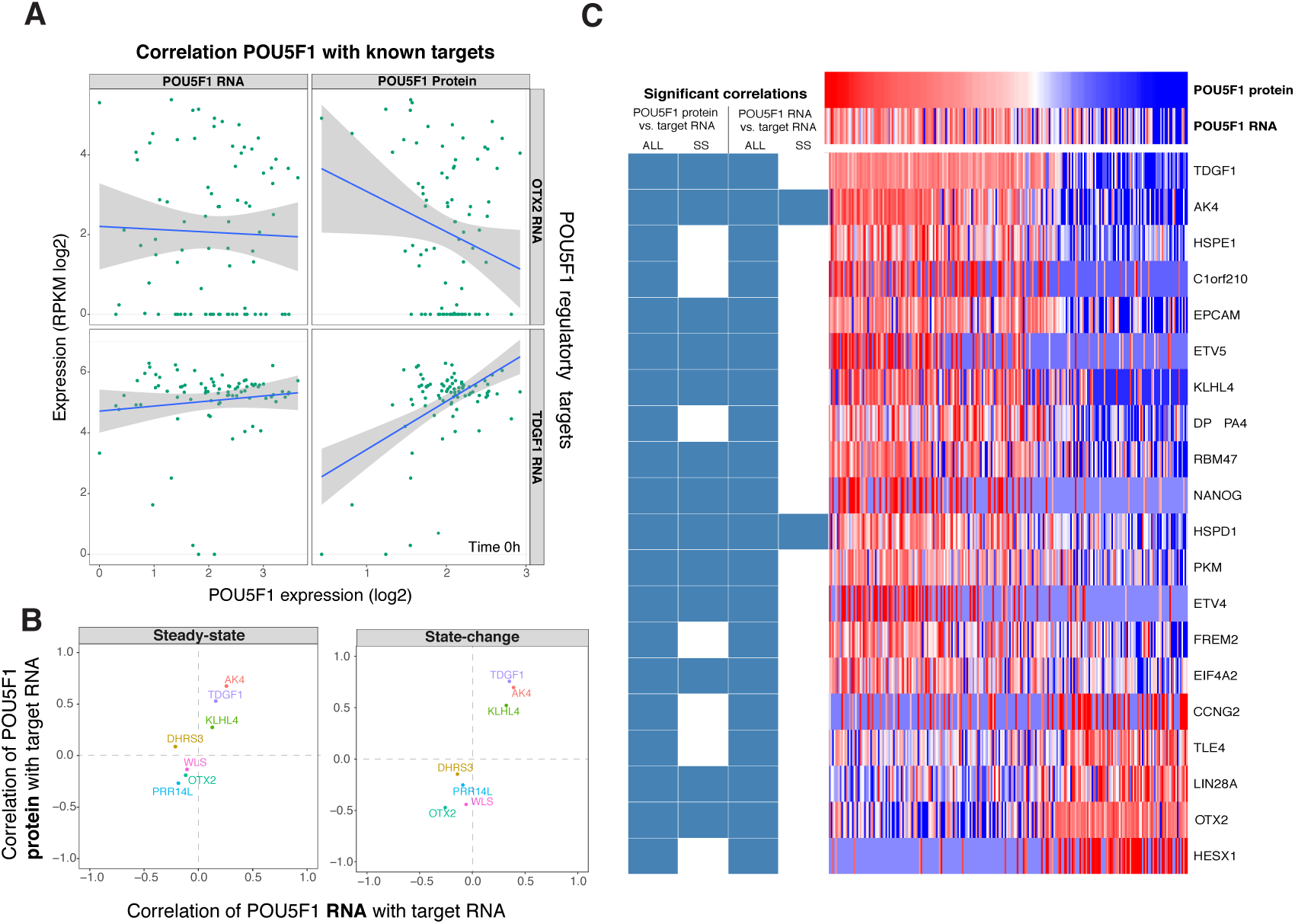
Transcription factor and trans regulatory correlation. **(A)** Scatter plot of POU5F1 expression in 0h single cells, both at the level of RNA (left panels) and protein (right panels) compared to the RNA expression of POU5F1 regulatory target genes OTX2 (upper panels) and TDGF1 (lower panels). Blue line represents linear model between the two expression patterns. **(B)** Scatter plot of Pearson correlation values of POU5F1_protein_-Target_RNA_ compared to Pearson correlation values of POU5F1_RNA_-Target_RNA_ for cells in steady state (0h) or undergoing a state-change (0h, 24h, and 48h). Each dot represents one target and the color of the dot identifies the gene. **(C)** Heatmap with expression levels of POU5F1_protein_, POU5F1_RNA_ and 20 genes_RNA_ identified as POU5F1 regulatory targets. The color scale relates to high expression (red) and low expression (blue). Blue rectangles indicate in which of the analysis groups the gene was considered to be a significant target. The ALL group includes samples from 0h, 24h and 48h. SS group includes samples from 0h or steady-state (SS) only. The columns in the heatmap are ordered based on the expression of POU5F1_protein_.

Given the strong correlation between levels of POU5F1_protein_ and a subset of known targets, we went on to investigate the possibility to use the correlation between protein expression of transcription factor and the expression of the specific transcripts as evidence that the corresponding gene might be regulated by that transcription factor. To test this, we ranked all TF-target scores (as described in Methods) of either POU5F1_protein_ or POU5F1_RNA_ as TF to all expressed genes in a set of samples. Next, we assigned all seven initial POU5F1-target pairs described above as positive targets and all the other genes as negative targets (n = 11,008). We used this classified ranked list to create a ROC curve to identify the cost of identifying negative targets, i.e. the false positive rate (FPR), based on the number of positive targets, i.e. true positive rate (TPR), identified. We did this using four different analysis groups where we calculated the correlation between POU5F1_protein or_ POU5F1_RNA_ and target_RNA_, and included either only steady-state (0h) cells or all cells (0h, 24h, 48h) **(Figure 3C)**.

To estimate the power of the different correlation analysis groups, we calculated the area under the curve (AUC) of the ROC curves. The results were as follows: correlation of POU5F1_protein_ and target_RNA_ with steady-state (AUC = 0.84) or all cells (AUC = 0.98), correlation of POU5F1_RNA_ and target_RNA_ with steady-state (AUC = 0.81) or all cells (AUC = 0.95) (**Figure S8)**. As expected, the AUC results suggest that using data from cells undergoing a state-change gives better power to detect positive POU5F1 targets than only using data from steady-state and that this is true for correlation to either POU5F1_protein_ or POU5F1_RNA_. Importantly, the results also show that POU5F1_protein_ expression levels better predict regulatory targets than POU5F1_RNA_ levels, especially when only using steady state samples.

We next asked if we could identify POU5F1 regulatory targets that were not in our initial candidate list. We did this by selecting the most correlated POU5F1-target pairs with a FPR of 0.10 for the four analysis groups analyzed above. Next, we filtered the list of gene pairs by requiring evidence that POU5F1 binds in the vicinity of the transcription start site of the identified target gene. To accomplish this, we used the same curated list (1035 genes) of published ChIP-seq data that we used for identifying the first seven candidates. This resulted in a list of 252 candidates where 73 genes were found in at least two of the analysis groups (**Figure 3C)**. Gene enrichment analysis of the 73 predicted POU5F1 targets using the Reactome pathway database identified two statistically significant (adjusted p-value < 0.05) pathways. Both of these pathways were directly related to the regulation of human pluripotent stem cells, specifically *Transcriptional regulation of pluripotent stem cells* (adjusted p-value = 0.0002) and *POU5F1, SOX2, NANOG activate genes related to proliferation* (adjusted p-value = 0.002). Also, POU5F1 was identified as the most significant transcription factor candidate in a query of the Enrichr TF-Gene Cooccurence data set (adjusted p-value 3.7e-14)^26^. In summary, our analysis highlights the power of using protein level measurements to identify transcription factor regulatory targets in single cell expression data, and also introduces an approach that provides orthogonal evidence to that from ChIP-seq for identifying target genes for transcription factors.

### Gene expression variation

A major strength of single-cell gene expression profiling is the possibility to study how gene expression varies between cells. Importantly, the factors that impact variation of expression at the mRNA or protein level can differ. To date, little is known about to what extent RNA expression variation translates into protein expression variation. For both mRNA and protein measurements taken at the 0h time point, we calculated the coefficient of variation of each gene. For mRNA, it is well understood that gene expression variation depends on the mean expression^27^, and the mRNA expression clearly shows this relationship **(Figure 4A).** We demonstrate a similar dependence for proteins **(Figure 4B).**

**Figure 4:**
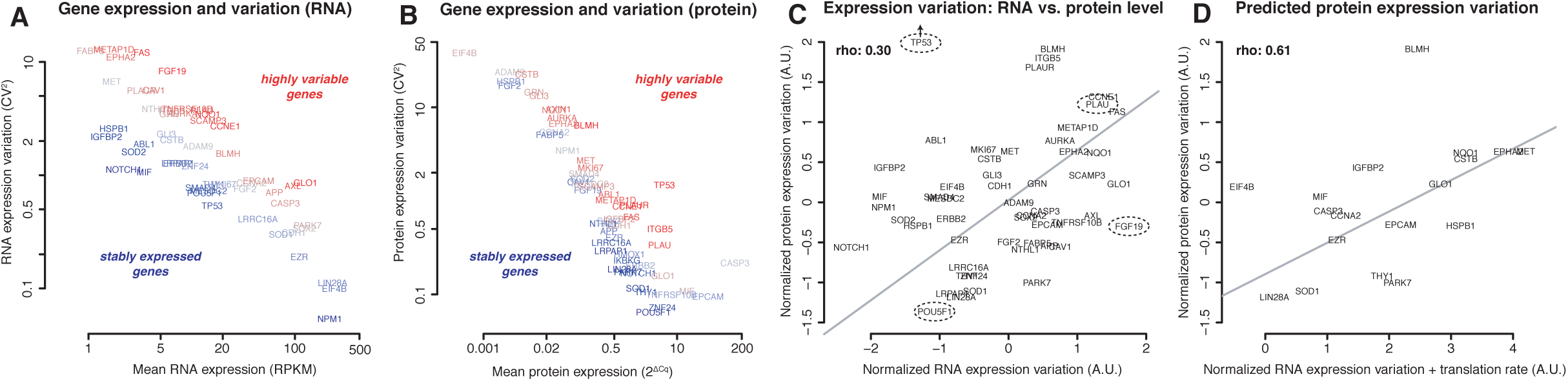
RNA and protein gene expression variation. Patterns of gene expression variation at steady-state in S-phase cells. **(A)** Relation between mean RNA expression mean and RNA expression variation (CV^2^). Colors indicate variable (red) or stable (blue) genes. **(B)** Similar to (A), but for protein. **(C)** Normalized gene expression variation (residuals) at the RNA and protein levels correlate moderately. Genes of interest are circled. TP53 has very high protein expression variation and is located outside the plot (x=−1.26, y=3.38). **(D)** A simple additive model of RNA variation and estimated translation rate effectively predicts protein expression variation (rho=0.61). Not all genes in Figure 3C are included here, since public data were not available to estimate translation rate for all genes.

We subsequently normalized the mRNA and protein expression variation data to their respective mean expression to obtain independent measurements of mRNA and protein expression variation (see Methods). We find that for many genes, variation of expression at the mRNA and protein level expression are generally comparable (Spearman’s rho=0.30, **Figure 4C**). The pluripotent factor POU5F1 is stably expressed both at the levels of RNA and protein, while other genes show both substantial RNA and protein expression variability (e.g. PLAU). Other subsets of genes are more variable on the mRNA level (e.g. FGF19), or on the protein level (e.g. TP53). Overall, these results highlight that variation at the mRNA level is not necessarily propagated to protein, and therefore RNA and protein levels should both be considered when assessing the impact of expression variation on gene function. Accordingly, SPARC provides valuable opportunities to study how mRNA variation relates to that at the protein level and how this process is regulated.

RNA and protein expression variation are complex, co-dependent processes. In order to investigate factors that determine differences between mRNA and protein expression variation, we considered the translation of mRNA to protein. We estimated translation rates by comparing mRNA abundances (RPKM) measured by SPARC with protein abundances from public quantitative mass-spectrometry data (see Methods). On the basis of reporter assays, it has been suggested that highly translated genes show higher variability of protein levels^28^. Here, we confirm this hypothesis for dozens of genes under physiological conditions as we find a positive correlation between the estimated translation rate and protein expression variability (Spearman’s rho=0.39). Interestingly, a simple addition of RNA expression variability and estimated translation rate yields an even better estimate of protein expression variability (Spearman’s rho=0.61, **Figure 4D**). This finding supports the notion that protein variability in single cells can be decomposed into RNA variability and noise originating from translation.

## DISCUSSION

We present a protocol, SPARC, to simultaneously profile mRNA and protein expression at cellular resolution. The modified Smart-seq2 protocol provides sensitive and precise gene expression detection and information of the full-length transcripts. Meanwhile, proximity extension assays enable highly specific and scalable protein detection of sets of cellular proteins in cells that have not been fixed. Using this method, we have explored the relationship between mRNA and protein expression in cells at steady-state or at the early stages of differentiation. Moreover, we introduce an approach to investigate mRNA and protein expression variation and covariance at cellular resolution, and extend the concept of mRNA velocity to use protein level expression data.

Overall, we find that the observed patterns of mRNA and protein co-expression reflect the snap-shot single cell measurements of gene expression with the inherent temporal element of delay between transcription and translation^6^ and the characteristics of mRNA and protein molecules, including expression kinetics^29^, half-lives, and copy numbers^2^.

Moreover, the observed relationships between mRNA and protein variability are consistent with earlier observations that cells employ mechanisms to either reduce or amplify intra- and inter-cell protein expression variability introduced by burst-like/noisy mRNA transcription^29-31^. As outlined by Raj and van Oudenaarden^31^, protein stability can play a role in the relationship between mRNA and protein variability. Short-lived proteins can track fluctuating mRNA levels closely, while levels of protein that degrade slowly fail to follow the rapid fluctuations of mRNA.

Going forward, measurement limitations and errors should be taken into consideration when evaluating mRNA-protein co-expression profiles. For example, no detection (zero expression) of a specific gene can arise either because its cognate mRNA was not present at the time the cell was interrogated (biological variation), or because the mRNA was lost during cDNA generation, amplification or library preparation (technical variation referred to as gene dropouts). While the Smart-seq2 procedure has high sensitivity and precision, technical variation is still present and needs to be considered when weighing the effects of biological versus technical variation^32^. For proteins, PEA assays have demonstrated low measurements variation^12^, they are less prone to dropouts but depend on antibody performance, and therefore may fail to detect some proteins despite relatively high protein expression.

The SPARC procedure described herein can help resolve regulatory networks or monitor developmental processes and cellular responses to e.g. genetic or chemical perturbations. We propose that the approach can be used for the analysis of transcript isoform usage and protein expression as explored in Gerlach et al^13^ and will be valuable for analyses of the relationship between mRNA and protein expression variation. The proximity-based protein assays are well suited to detect post-translational modifications^33^ and can be scaled up via a sequencing readout^34^, allowing measurement of still greater numbers of proteins and protein modifications in individual cells.

In summary, the more direct roles of proteins in maintaining cellular functions compared to transcripts, together with the recognized importance of posttranslational regulation, render single cell protein analysis an important complement of comprehensive RNA analyses of cell state and to decipher gene regulation circuitry.

## MATERIAL AND METHODS

### Cell culture and neural induction of human embryonic stem cells

Human embryonic stem cell line HS181 (hPSCreg Kle001-A) was maintained on vitronectin (VTN-XL; Stem Cell Technologies, 07180) in Essential-E8 Medium (ThermoFisher Scientific, A1517001). Cells were passaged as clumps with gentle cell dissociation reagent (GCDR; Stem Cell Technologies, 07174) when necessary. Two days before neural induction, cells were harvested with GCDR and plated in Essential-E8 supplemented with 10 uM Rho-kinase inhibitor Y27632 (Stem Cell Technologies, 72304) on VTN coated 6-well culture dish to yield approximately 80% confluence next day. When cells reached 100% confluence, neural differentiation was induced (Day 0/0h) using dual smad inhibition (Maroof et al., 2013). Briefly, cells were washed with 1x DPBS and neural induction medium (NIM) was added, consisting of KnockoutTM DMEM (10829018), 15% KnockOutTM Serum Replacement (A3181501), 1x GlutaMax (35050038), 1x non-essential amino acids (11140035), 1% penicillin/streptomycin (1514022)(all ThermoFisher Scientific), supplemented with 2 uM tankyrase inhibitor XAV939 (Sigma-Aldrich, X3004), 100 nM ALK2/3 inhibitor LDN193189 (Miltenyi Biotech, 130-106-540) and 10 uM ALK4/5/7 inhibitor SB431542 (Millipore, 616464). Medium was replaced on days 1 and 2.

To follow early neural differentiation, cells were harvested at indicated time points (0h; 24h; 48h). Cells were washed twice with DPBS to remove floating dead cells, and subsequently treated with Accutase (Sigma-Aldrich) for 5 to 15 min until a single cell suspension was obtained. Cells were collected in 13 ml DPBS and then centrifuged at 300 × g for 5 min, washed with DPBS, centrifuged again and re-suspended in DPBS. Cells were kept on ice until single cell sorting.

### Single cell isolation

The single cells were incubated on ice with a viability dye (LIVE/DEAD Viability Kit, Molecular Probes, L3224) for approximately 15 minutes and then filtered through a 40 μM filter before sorting. Cells were sorted on a BD FACS ARIAIII into a 96-well plate. We used a gating strategy to exclude of debris, doublets and cells positive for EthD-1, a nucleic acid dye that enters cells with damaged membranes. After the cells were sorted, the plate was centrifuged at 700 g and 4°C for 1 minute and then quickly transferred to dry ice. The plates were stored at −80°C until processed.

For the SPARC protocol, cells were sorted into 1.5 μl TE buffer (pH 8.0 Invitrogen, AM9858) with the following components: 1% NP-40 (Thermo Fisher, 28324), 0.1% Triton X-100 (Thermo Fisher, 28314), 0.1% Sulfobetaine (Sigma-Aldrich, 82804-50G), 150 mM NaCl (Ambion, AM9760G), 10 mg/ml BSA (Ambion, AM2616), 2 U SUPERase In RNase Inhibitor (Ambion, AM2696), 1X HALT protease inhibitor (Thermo Fisher, 78430) and 1:1,250,000 ERCC (Invitrogen, 4456740).

For the Smart-seq2 protocol, cells were sorted into 4 μl TE buffer pH 8.0 with the following components: 0.1% Triton X-100 (Thermo Fisher, 28314), 1 U SUPERase In RNase Inhibitor (Ambion, AM2696), 1:4000000 ERCC spike-in (Ambion, 4456740), 2.5 mM dNTPs (Thermo Fisher, R0191) and 2.5 μM oligo-dT (Integrated DNA Technologies IDT, 5’-/5BiotinTEG/AAGCAGTGGTATCAACGCAGAGTA CT_30_VN-3’, where V is either A, C or G, and N is any base). For multiplex protein detection only, cells were sorted into 1.5 μl TE buffer pH 8.0 with the following components: 1% NP-40 (Thermo Fisher), 0.1% Triton X-100 (Thermo Fisher), 0.1% Sulfobetaine (Sigma-Aldrich), 150 mM NaCl, 10 mg/ml BSA (Ambion, 4456740), and 1X HALT Protease Inhibitor (Thermo Fisher).

On each 96-well sorting plate, we included: (i) population controls of 100 sorted cells/well in duplicate; (ii) buffer, no-cell control in triplicate; (iii) 100 cell equivalent lysate prepared from the SK-MEL-30 cell line. The SK-MEL-30 cell lysate was prepared in bulk, aliquoted and added to every plate in triplicate as an inter-plate control.

### Oligo-dT bead preparation

For every reaction, 5µl Dynabeads MyOne Streptavidin T1 (Invitrogen, 65602) was washed twice with 1.45 µl washing solution containing 100mM NaOH (Sigma-Aldrich, S8045-500G), 50mM NaCl (Ambion, AM9760G) and UltraPure DNase/RNase-Free Distilled Water (Invitrogen, 10977035). The beads were then washed with 1.45 µl RNAse-free bind and wash solution containing 0.01 mM Tris (Invitrogen, AM9855G), 1mM EDTA (AM9260G), 2 M NaCl (Ambion, AM9760G) and UltraPure DNase/RNase-Free Distilled Water (Invitrogen, 10977035). The beads were then mixed with 2 µl RNAse-free bind and wash solution and 0.1 µl oligo-dT (IDT, 5’-/5BiotinTEG/AAGCAGTGGTATCAACGCAGAGTA CT_30_VN-3’) and incubated in ambient temperature for 15 minutes. The beads were finally stored in 1 µl 1% BSA (Ambion, AM2616) in TE buffer (Invitrogen, AM9858) and incubated on a rotator overnight in 8°C. Before use, the buffer was exchanged for the RNA and Protein lysis buffer.

### mRNA capture, reverse transcription and pre-amplification

Lysis plates with FACS sorted cells were centrifuged in 700 g for 10 seconds and thawed on ice. Smart-seq2 reference samples were incubated at 72°C for 3 minutes and directly placed back on ice. SPARC samples were pipette mixed while 1 µl of prepared beads was added to each reaction. The SPARC plate was then incubated for 10 minutes with orbital shaking at 1000 rpm. Plates were then centrifuged at 700 g for 10 seconds and placed on a magnetic rack (Alpaqua Magnum FLX) where 1.7 µl from each well was transferred for protein analysis. The Smart-seq2 and SPARC samples were supplied with 6 µl and 10 µl reverse transcription mix, respectively. The mixes contained: 100 U SuperScript II reverse transcriptase, 1x First Strand Buffer, 5mM DTT (all Invitrogen, 18064014), 10 U SUPERase In RNase Inhibitor (Invitrogen, AM2696), 1 M Betaine (Sigma-Aldrich, 61962-250G), 6 mM MgCl_2_ (Invitrogen, AM9530G), 1 mM each of dNTP’s (ThermoScientific, R0192), 1 µM TSO (5’- AAGCAGTGGTATCAACGCAGAGTACATrGrG+G-3’, Exiqon as described in (Picelli et al., 2014) and UltraPure DNase/RNase-Free Distilled Water (Invitrogen, 10977035). The Smart-seq2 samples were centrifuged for 700 g for 10 seconds and then placed in a thermal cycler. The SPARC samples were pipette mixed prior to incubation in a thermal cycler. Both samples sets were processed using the following program: 42°C for 90 minutes, 10 cycles with 50°C for 2 minutes and 42°C for 2 minutes and finally 70°C for 15 minutes before holding at 4°C.

Each first strand cDNA reaction was supplied with 15 µl of PCR mix containing 1x KAPA HiFi HotStart Ready Mix (Kapa Biosystems, KK2601), 1 µM IS PCR primer (5’-AAGCAGTGGTATCAACGCAGAGT-3’, IDT, as described in (Picelli et al., 2014) and UltraPure DNase/RNase-Free Distilled Water (Invitrogen, 10977035). The RNA samples were vortexed and centrifuged (700 g for 10 seconds) while the combined RNA/Protein samples were pipette mixed, prior to incubation in PCR program: 98°C for 3 minutes, cycling of 98°C for 20 seconds, 67°C for 15 seconds and 72°C for 6 minutes. Lastly, the final extension was at 72°C for 30 seconds prior to holding at 4°C. Single cells were subjected to 20 PCR cycles while bulk samples (100 cells) were subjected to 14 PCR cycles. Samples were finally purified with AMPure XP beads (Beckman Coulter, A63880) using 0.8X bead to sample ratio.

### scProtein expression analysis

Cell lysate containing the protein supernatant were transferred to a new 96-well PCR-plate and were processed immediately. To each sample, we added 2.1 µl Incubation Solution (Olink Proteomics), 0.3 µl Incubation Stabilizer (Olink Proteomics), 0.3 µl of each PEA A- and B-probe mix (final concentration 100 pM; Olink Proteomics). The probes targeted 92 cellular proteins and 4 controls. The controls included spiked-in GFP, PE, an extension control and a detection control^15^. Each 96-well plate included a lysis buffer only negative control in triplicate. PCR-plates were briefly vortexed, centrifuged, sealed and incubated overnight at 8°C. Following overnight incubation, plates were brought to room temperature, briefly spun down and 96 µl Extension mix was added to each well. The extension mix contained 10 µl PEA Solution (Olink Proteomics), 0.5 µl PEA Enzyme (Olink Proteomics), 0.2 µl PCR Polymerase (Olink Proteomics) and 85.3 µl UltraPure DNase/RNase-Free Distilled Water (Invitrogen, 10977035). Plates were sealed, gently vortexed, centrifuged and within 5 minutes of adding the Extension mix, placed in a thermal cycler for the extended reaction (50 °C, 20 min), and pre-amplification of extended PEA probes via universal primers (95°C, 5 min, (95°C, 30 s; 54°C, 1 min; and 60°C, 1 min) × 17).

The pre-amplified extended PEA products were decoded and quantified using a Fluidigm 96.96 Dynamic Array Integrated Fluidic Circuit on a Biomark HD system. Ninety-six primers pairs (5 µl of each) targeting each PEA probe pair were loaded in the left inlets of the array. A Detection mix containing 5 µl Detection Solution, 0.071 µl Detection Enzyme, 0.028 µl PCR Polymerase (all Olink Proteomics) and 2.1 µl UltraPure DNase/RNase-Free Distilled Water (Invitrogen, 10977035) was added in the right inlets of the array. The 96.96 IFC chip was primed in Fluidigm’s IFC HX according to manufacturer’s instructions and then run on the Biomark HD system with the following settings: Gene Expression application, ROX passive reference, single-probe assay with FAM-MGB probe. The thermal protocol included thermal mix (50C, 120 s; 70C, 1,800 s; 25C, 600 s), hot start (95C, 300 s), and PCR cycling for 40 cycles (95C, 15 s; 60C, 60 s).

### Library preparation and sequencing

Purified cDNA (75 ng) was used as input to the Nextera XT DNA library preparation kit (Illumina, FC-131- 1096), following the manufacturers protocol with the modification of using 1:5 of reagent volumes. Indexing primers (Illumina, FC-131-2001; FC-131-2002; FC-131-2003; FC-131-2004) were diluted 1:2 in UltraPure DNase/RNase-Free Distilled Water (Invitrogen, 10977035) prior to use. Samples were finally pooled and purified with AMPure XP beads (Beckman Coulter, A63880) using 0.6x bead to sample ratio and concentrated by eluting in 60% of the corresponding input sample volume using Elution buffer (Qiagen, 19086). The final sequencing library was quantified using BioAnalyzer High Sensitivity DNA kit (Agilent, 5067-4626) using a region table spanning 100 bp to 1000 bp. The pooled library was sequenced on two lanes of Illumina HiSeq2500 using single read 50 bp read length, v4 chemistry. The sequencing was performed at the SNP&SEQ Technology Platform, Science for Life Laboratory, Uppsala, Sweden.

### scRNAseq dataset processing

Reads were mapped to the human genome (GRCh38) including the sequence of the spike-in RNAs. FeatureCount summarised over annotated genes from the GRCh38.77 version of the human genome including the spike ins were used to get counts for all exons of annotated genes in the human genome. Samples with less than 10 000 reads mapping to the exons or the fraction of spike-in RNAs were greater than 20 percent were removed from further analysis. RPM values and RPKM values were calculated for all genes in all samples.

### scProtein dataset processing

Following the completion of the qPCR run on the Fluidigm Biomark HD system, we visually inspected the amplification curves. Samples showing evidence of failed or poor amplification reactions were excluded from further analysis. Next, the raw Cq data (log 2 scale) from the Fluidigm Biomark HD system was exported and processed. First, samples were excluded if no signal was detected in any of control assays (extension control, incubation control or detection control), or if the signal in any of the controls was greater or less than 2 standard deviations (SD) of the mean value across all samples measured on a 96.96 IFC Biomark chip. Next, the remaining Cq values were normalized for intra-plate variation with the extension control (Cq_assay_ – Cq_ExtCtl_) yielding dCq values. Then, for each assay, the dCq values were subtracted from the negative control computed as the lysis buffer mean + 2*SD. This ensures that observed signals for each assay in the presence of a cell are at least 2 SDs away from any signals observed in the absence of any antigen. Resulting values below zero were set to zero and the signal was deemed undetected. The cumulative protein sum was calculated by summing across all proteins measured (n=92) per cell.

### Method comparison for RNA analysis

Samples from both the SPARC protocol and standard Smart-seq2 protocol at 0h were used to compare similarities and differences between the two protocols. Only genes with log RPKM greater than 1 were used for further analysis. Genes were separated into different biotypes according to GRCh38.77 annotation and number of detected genes per biotype were compared between the two protocols. Read counts for all exons and introns in all genes was counted and the distribution across the genes were compared between the two protocols. Logarithmic mean expression (LME) for each gene was calculated for all samples irrespective of protocol, and for each protocol by itself LMESPARC-seq and LMESmart-seq2. Differential expression between the protocols (DE-prot) per gene were calculated by dividing the LMESPARC-seq with the LMESmart-seq2 value per gene. Both the LME and the DE-prot was taken into account by multiplying the two values to identify the differences between the two protocols. Genes were the product of the two was greater than 8 was considered to be different and analyzed for differences in lengths and gene biotype.

### Cell cycle assignment

RNA samples were normalized using the Seurat 2^24^ package and the samples were scaled by the number of detected genes per sample. Scores for each sample being in either S phase or G2/M phase was calculated using the Seurat 2 package and at the same time predicted to belong to either the G1,S or G2/M phase.

To further explore the relationship between cell cycle phase and protein expression, the hESCs were labeled with the Live cell DNA dye Vybrant DyeCycle Violet (Invitrogen, V35003) and sorted by cell cycle phase (G1, S or G2/M) in triplicate at 100 cells per well. Cells were sorted in the following lysis buffer: 2 μl TE buffer pH 8.0 with the following components: 1% NP-40 (Thermo Fisher, 28324), 0.1% Triton X-100 (Thermo Fisher, 28314), 0.1% Sulfobetaine (Sigma-Aldrich), 150 mM NaCl (Thermo Fisher, AM9760G), 10 mg/ml BSA (Thermo Fisher, AM2618), and 1X HALT Protease Inhibitor (Thermo Fisher, 78430). Cells were then processed for multiplex PEA analysis as described above.

### PCA analysis and pseudotime analysis

The normalized and scaled data from the cell cycle prediction were further scaled by the scores for the S phase and the G2M scores to remove the cell cycle dependency of the samples using the Seurat package. Dimensional reduction analysis using tSNE were used to reduce the dimensionality of the sample data. The three dimensions from the tSNE analysis was then used by SCORPIUS^19^ to predict a linear pseudo time through all the samples with a score between 0 and 1. Average pseudotime scores for the 0h samples and 48h samples were calculated. If the average pseudotime score for the 0h samples were higher than the average 48h samples the pseudotime was re-calculated by subtracting the pseudotime with 1 and then multiplying by −1 to change the order of the samples and maintaining the pseudotime distances between samples. Pseudotime scores were then multiplied with 48 to reflect a pseudotime over 48h. Genes that change over time was identified by SCORPIUS (adj p-value <0.05) and separated into different modules dependent on expression pattern over time.

### Comparison of RNA and protein changes over time

To test whether the expression levels for both RNA and protein level show the same directional changes, we applied a linear model for expression over time for both the RNA logged RPKM values (lRPKM(pt) = klRPKM*pt + IlRPKM) and for protein Cq values (Cq(pt) = kCq*pt + ICq). We then tested if the slope of the RNA levels for a gene could predict the slope of the protein for the same gene with the linear model (lRPKM(pt) = klRPKM*pt + IlRPKM) (Cq(pt) = kCq*pt + ICq). We then tested if the slope of the RNA levels for a gene could predict the slope of the protein for the same gene with the linear model kCq(klRPKM) = klRPKM*0.49 + 0.01.

### RNA and protein velocity

Exon, intron and spanning reads were retrieved using the reads mapped to the GRCh38 reference and the GRCh38.77 annotation. Only coding genes were the total intron counts and exon counts were greater than 1000 reads and spanning gene count reads with more than 500 reads were kept for further analysis. To estimate velocity we used 20 cell kNN pooling with the gamma fit based on an extreme quantile of 0.05 including the spanning reads the gene offsets. The flow of the cells was visualised on the tSNE embedding that was identified using the Seurat 2 package described above and used in the pseudotime analysis. To generate a set of noise samples, intron counts per gene per sample were randomly shuffled between samples to remove true correlation between exon and introns. The same parameters and method that was used to get the true flow was used for the shuffled exon intron dataset. For the mature mRNA and protein analysis, RPKM values for exons as “intron” counts and Cq values for protein were used as “exon” counts. All 62 proteins that passed protein QC was used for analysis. To estimate the velocity, we used 20 cell kNN pooling with the gamma fit based on an extreme quantile of 0.05. The flow of the cells were visualised on the tSNE embedding that was identified using the Seurat 2 package^24^ described above and used in the pseudotime analysis. As a further control, the assignment of gene RNA expression values and gene protein expression values was permuted for each cell, thereby preserving the total RNA and protein abundance in each cell.

### Gene regulatory network analysis – selection of POU5F1 targets

To identify genes that are trans-regulated by POU5F1, an initial set of targets to POU5F1 was selected based on two criteria: (1) It should be identified as changing over pseudotime according to the pseudotime calculated and described above; and (2) It should be reported as bound by POU5F1 in the vicinity of the TSS by the curated set of ChIP-seq data described in the section below. The intersection of the two assumptions gave a subset of 7 genes.

A set of genes where POU5F1 binds in the vicinity of the TSS in primed hESCs were identified using ChIP-seq data^35^. An initial set of 11 hESC samples (SRX017276, SRX021069, SRX021071, SRX1053369, SRX1053370, SRX1053378, SRX1053379, SRX266859, SRX702065, SRX702066, SRX702069) were collected from CHiP-Atlas^35^ by filtering with POU5F1 as antigen and a distance from TSS as 1 kb. For the remaining samples hierarchical clustering of the euclidean distance all samples binding scores across all genes compared to each other was done. A final set of 7 samples (SRX017276, SRX021069, SRX021071, SRX1053378, SRX1053379, SRX702065, SRX702066) that clustered as primed hESC in the hierarchical cluster were kept for further analysis. Genes with a reported binding score in ChIP-Atlas in at least two samples were considered as the curated set of POU5F1 target genes.

TF-target scores were calculated for a set of sc samples in three steps: (1) calculation of the Pearson correlation between the TF expression pattern and the expression pattern of annotated genes in a cell; (2) calculation of a z-score for each TF-target by subtracting the mean and dividing by the standard deviation of the Pearson correlation scores for all TF-targets; and (3) The TF-targets scores was then calculated by taking the absolute value of the z-scores creating a distribution scores greater than zero. The greater the TF-targets score the more correlation, positive or negative, between the TF and the target expression pattern.

### Gene expression variation

Gene expression variation was estimated using the squared coefficient of variation (CV^2^). As variation and mean expression are inherently linked due to sampling properties, we perform a linear fit of variation to mean expression in log-space. For protein measurements, a cubic polynomial was applied to reflect the non-linear dependence of exceedingly lowly or highly expressed proteins. For RNA measurements, weights equal to the square root of the expression values were applied to the fit to withhold lowly expressed genes from driving the fit. We then consider the residuals of the individual gene measurements with regard to this fit as mean independent gene expression variation measurements (termed “normalized variation”). To avoid in-silico biases, data was not computationally de-noised using batch-effect removal tools. Instead, cells were strictly filtered and quality controlled. Specifically, outliers were removed using pseudo-time estimations from SCORPIUS, and only cells labeled S-phase of the cell-cycle using the cyclone package of SCRAN were considered. Mass-spectrometry data for the estimation of the translation rate for hESCs (E14 cells^36^) was retrieved from the FunCoup PaxDB statistics^37^. Translation rates are estimated as the log2 ratio of protein abundances vs. RNA RPKM measurements. Due to the different dynamic ranges of the RNA variability and translation measures, the “combined normalized variation and translation rate” reflects the sum of normalized RNA expression variation and 0.5 times estimated translation rate. Lines in Figures 4C and D show linear least total square fits.

## Data availability

The RNA expression data is uploaded to ENA with the project number PRJEB33157. The protein data is attached as **Table S2.**

## SUPPLEMENTAL RESULTS & FIGURES

### Relation of RNA and protein data in single cells at steady-state

We observed a number of genes where mRNA and protein expression are discordant. Specifically, we observed a very low fraction of cells expressing mRNA but a high fraction of cells expressing the cognate protein. A parsimonious explanation for these observations is that we failed to reliably capture, amplify and sequence the specific mRNAs, an outcome that is common in scRNAseq experiments (ref). To explore this possibility, we isolated total RNA from the same hESC cell line used in the study and performed bulk cell gene expression analysis using TaqMan Gene Expression assays. We targeted three genes showing discordant gene expression (IKBKG, HMOX1, METAP1D) and included EIF4B as a positive control. We clearly detect RNA expression in all the genes analyzed, suggesting that the mRNA for IKBKG, HMOX1, METAP1D are expressed but were not reliably detected in the single cell experiments.

However, we cannot rule out that the observed discordant mRNA and protein co-expression at the single cell level is due to that the mRNAs for these genes have very short-half lives and are absent in many cells, whereas the cognate proteins are relatively stable. Alternatively, but less likely, the protein expression signal originates from antibody cross-reactivity to homologous gene products. PEA probes employed in this study were tested to ensure that they indeed detect the expected target, and that they do not give any signal when tested against a large pool of recombinant proteins.

### TaqMan Gene Expression analysis

Total RNA was isolated from the HS181 cell line using the miRNeasy Micro Kit (Qiagen, 217084). We used the One-Step RT-PCR System (Thermo Fisher Scientific, 12574026) with the Taqman Assays IKBKG (Hs00415849_m1, 4453320), METAP1D (Hs00994998_m1, 4448892), HMOX1 (Hs01110250_m1, 4453320) and EIF4B (Hs00973573_m1, 4331182) (all Thermo Fisher). Gene expression was quantified from the isolated RNA at 100 ng/reaction using the Quantstudio real time qPCR instrument.

**Figure S1: Characteristics of SPARC**

**(A)** Comparative characteristics of the single cell mapped sequencing reads (exon, intergenic, intron) for the SPARC versus Smart-seq2 protocol. Data is reported both for the reads originating from the single cell data and the FACS sorted 100 cell control data. **(B)** Annotation of differentially expressed genes detected via SPARC versus Smart-seq2. The genes preferentially detected with SPARC tend to be longer than the genes preferentially detected with Smart-seq2. The Not DE group represent the group of genes equally detected with both methods. **(C)** Single cell mean RKPM (log2) expression for all genes measured using either SPARC (x-axis) or Smart-seq2 (y-axis) (Pearson correlation coefficient = 0.90) **(D)** Comparison of SPARC mean RKPM (log2) RNA expression reads for replicate FACS sorted 100 hESC cells at 0h population (x-axis) or single cells (Pearson correlation coefficient = 0.96). **(E)** Comparison of Smart-seq2 mean RKPM (log2) RNA expression reads for replicate FACS sorted 100 hESC cells at 0h population (x-axis) or single cells (Pearson correlation coefficient = 0.96).

**Figure S2: RNA and protein gene expression violin plots**

Gene expression plots for the genes measured at both the level of protein (red) and RNA (blue). Results are shown for both the duplicate 100 cell control (black dots) and single cells (violin plots). The presented data is not normalized for cell cycle, the number of genes detected (RNA) or cumulative protein sums (protein).

**Figure S3: Cell cycle and gene expression**

Relationship between the predicted cell cycle phase (G1, S or G2/M) and **(A)** the per cell cumulative protein sum or **(B)** number of detected genes. The cumulative protein sum was calculated by summing across all proteins measured (n=92) per cell. **(C)** hESCs were labeled with the live cell Vybrant DyeCycle Violet DNA stain and sorted by cell cycle phase G1, S or G2/M. Cells were sorted 100 cells in triplicate per cell cycle and processed for multiplex PEA analysis. Protein expression was compared in G2/M sorted cells compared to G1 sorted cells. The volcano plot shows the extent of the difference in expression between G2 and G1 cell cycle phases (x-axis) and significance of the difference (y-axis). The red line marks p-value 0.01. The blue lines mark 0.5 Cq difference between the cell cycle phases G2 and G1, or an approximate 1.5-fold difference. As expected, G2-specific proteins AURKA, CCNA2 and AURKB and G1-Gspecific protein CCNE1 show higher expression in G2 and G1 cell cycle phases, respectively. The majority of protein show a mean 1.5 increase in protein amount in G2 versus G1, with some notable exceptions, including e.g. pluripotent factors NANOG and POU5F1.

**Figure S4: RNA and protein gene expression dot and density plots**

Combined dot and density plot of mRNA (log RPKM) and protein (Cq) expression in cells measured at 0h (green), 24h (orange) and 48h (blue). The presented data is not normalized for cell cycle, the number of genes detected (RNA) or cumulative protein sums (protein).

**Figure S5: RNA and protein gene expression in single cells as ordered by pseudotime**

mRNA (log RPKM) and protein (Cq) expression in cells ordered by pseudotime as determined by the RNA expression data. The presented data is not normalized for cell cycle, the number of genes detected (RNA) or cumulative protein sums (protein).

**Figure S6: Analysis of agreement of mRNA or protein expression changes over time**

Agreement of changes of mRNA and protein abundances over the measured time points 0h, 24h and 48h at the level of RNA and protein. The plot shows the results of the linear model Cq(lRPKM)) = 0.49* lRPKM + 0.01 with a significant Pearson r-square value of 0.20 (p-value 5,1*10^−4^). Each dot represents a measured gene.

**Figure S7: RNA and protein velocity controls**

**(A)** The results of the RNA velocity analysis. **(B)** Example result of one round where we permuted the intron count per gene between samples to test if sample embedding within the tSNE plot can influence the direction of the velocity vectors. Color corresponds to time point in all plots 0h (green), 24h (orange) and 48h (blue).

**Figure S8: ROC-curve of Pearson correlation scores as predictor of POU5F1 targets**

Red lines represent POU5F1_protein_-target_RNA_ ROC curves and blue lines represent POU5F1_RNA_-target_RNA_ ROC curves. Solid line represents when all samples (0h, 24h, 48h) have been considered and dashed lines represents when only steady-state (0h) samples have been considered. Filled circles represent the Youden index of the different ROC curves.

**Table S1: List of the PEA protein expression assays**

**Table S2: Protein expression data**

0h, 24h and 48h protein expression data for the single cell and 100 cell FACS controls. The data is reported as described in methods under scProtein dataset processing but are not normalized for cumulative protein sum. Assays are filtred for those where we detected the protein at a level of > 3 Cq over background in the 100 cell population control in at least one time point.

## ACKNOWLEDGEMENTS

CG acknowledges funding from the Swedish Research Council (VR) grant 2017-05229 and support from the Single Cell Proteomics Facility, Science for Life Laboratory, Sweden. MT and MRF acknowledge funding from ERC Starting Grant 758397 and from the Strategic Research Area (SFO) program of the Swedish Research Council through Stockholm University. ND acknowledges funding from the Swedish Research Council 2015-02424. Sequencing was performed by the SNP&SEQ Technology Platform in Uppsala, Sweden. The facility is part of the National Genomics Infrastructure (NGI) Sweden and Science for Life Laboratory. The SNP&SEQ Platform is also supported by the Swedish Research Council and the Knut and Alice Wallenberg Foundation. We thank Ulf Landegren and Lars Feuk for helpful feedback on the manuscript.

## REFERENCES

1. Larsson, A.J.M. et al. Genomic encoding of transcriptional burst kinetics. Nature 565, 251–254 (2019).

2. Schwanhäusser, B. et al. Global quantification of mammalian gene expression control. Nature 473, 337–342 (2011).

3. Natarajan, K.N. et al. Comparative analysis of sequencing technologies for single-cell transcriptomics. Genome Biol 20, 70 (2019).

4. Jovanovic, M. et al. Immunogenetics. Dynamic profiling of the protein life cycle in response to pathogens. Science 347, 1259038 (2015).

5. Vogel, C. & Marcotte, E.M. Insights into the regulation of protein abundance from proteomic and transcriptomic analyses. Nature reviews. Genetics 13, 227–232 (2012).

6. Liu, Y., Beyer, A. & Aebersold, R. On the Dependency of Cellular Protein Levels on mRNA Abundance. Cell 165, 535–550 (2016).

7. Peterson, V.M. et al. Multiplexed quantification of proteins and transcripts in single cells. Nat Biotechnol 35, 936–939 (2017).

8. Stoeckius, M. et al. Simultaneous epitope and transcriptome measurement in single cells. Nat Methods 14, 865–868 (2017).

9. Popovic, D., Koch, B., Kueblbeck, M., Ellenberg, J. & Pelkmans, L. Multivariate Control of Transcript to Protein Variability in Single Mammalian Cells. Cell Syst 7, 398–411 e396 (2018).

10. Frei, A.P. et al. Highly multiplexed simultaneous detection of RNAs and proteins in single cells. Nat Methods 13, 269–275 (2016).

11. Schulz, D. et al. Simultaneous Multiplexed Imaging of mRNA and Proteins with Subcellular Resolution in Breast Cancer Tissue Samples by Mass Cytometry. Cell Syst 6, 531 (2018).

12. Darmanis, S. et al. Simultaneous Multiplexed Measurement of RNA and Proteins in Single Cells. Cell reports 14, 380–389 (2016).

13. Gerlach, J.P. et al. Combined quantification of intracellular (phospho-)proteins and transcriptomics from fixed single cells. Sci Rep 9, 1469 (2019).

14. Picelli, S. et al. Smart-seq2 for sensitive full-length transcriptome profiling in single cells. Nat Methods 10, 1096–1098 (2013).

15. Assarsson, E. et al. Homogenous 96-plex PEA immunoassay exhibiting high sensitivity, specificity, and excellent scalability. PloS one 9, e95192 (2014).

16. La Manno, G. et al. RNA velocity of single cells. Nature 560, 494–498 (2018).

17. Schmid, M. & Jensen, T.H. Controlling nuclear RNA levels. Nature reviews. Genetics 19, 518–529 (2018).

18. Padovan-Merhar, O. et al. Single mammalian cells compensate for differences in cellular volume and DNA copy number through independent global transcriptional mechanisms. Mol Cell 58, 339–352 (2015).

19. Cannoodt, R. et al. SCORPIUS improves trajectory inference and identifies novel modules in dendritic cell development. bioRxiv (2016).

20. Liu, Y. & Aebersold, R. The interdependence of transcript and protein abundance: new data--new complexities. Mol Syst Biol 12, 856 (2016).

21. Ziller, M.J. et al. Dissecting neural differentiation regulatory networks through epigenetic footprinting. Nature 518, 355–359 (2015).

22. Lee, M.V. et al. A dynamic model of proteome changes reveals new roles for transcript alteration in yeast. Mol Syst Biol 7, 514 (2011).

23. Gaidatzis, D., Burger, L., Florescu, M. & Stadler, M.B. Analysis of intronic and exonic reads in RNA-seq data characterizes transcriptional and post-transcriptional regulation. Nat Biotechnol 33, 722–729 (2015).

24. Butler, A., Hoffman, P., Smibert, P., Papalexi, E. & Satija, R. Integrating single-cell transcriptomic data across different conditions, technologies, and species. Nat Biotechnol 36, 411–420 (2018).

25. Qiu, X. et al. Towards inferring causal gene regulatory networks from single cell expression Measurements. bioRxiv, 426981 (2018).

26. Kuleshov, M.V. et al. Enrichr: a comprehensive gene set enrichment analysis web server 2016 update. Nucleic Acids Res 44, W90–97 (2016).

27. Grun, D. & van Oudenaarden, A. Design and Analysis of Single-Cell Sequencing Experiments. Cell 163, 799–810 (2015).

28. Ozbudak, E.M., Thattai, M., Kurtser, I., Grossman, A.D. & van Oudenaarden, A. Regulation of noise in the expression of a single gene. Nat Genet 31, 69–73 (2002).

29. Raj, A., Peskin, C.S., Tranchina, D., Vargas, D.Y. & Tyagi, S. Stochastic mRNA synthesis in mammalian cells. PLoS Biol 4, e309 (2006).

30. Bahar Halpern, K. et al. Nuclear Retention of mRNA in Mammalian Tissues. Cell reports 13, 2653–2662 (2015).

31. Raj, A. & van Oudenaarden, A. Nature, nurture, or chance: stochastic gene expression and its consequences. Cell 135, 216–226 (2008).

32. Ziegenhain, C. et al. Comparative Analysis of Single-Cell RNA Sequencing Methods. Mol Cell 65, 631–643 e634 (2017).

33. Weibrecht, I. et al. In situ detection of individual mRNA molecules and protein complexes or post-translational modifications using padlock probes combined with the in situ proximity ligation assay. Nat Protoc 8, 355–372 (2013).

34. Darmanis, S. et al. ProteinSeq: high-performance proteomic analyses by proximity ligation and next generation sequencing. PloS one 6, e25583 (2011).

35. Oki, S. et al. ChIP-Atlas: a data-mining suite powered by full integration of public ChIP-seq data. EMBO Rep 19(2018).

36. Phanstiel, D.H. et al. Proteomic and phosphoproteomic comparison of human ES and iPS cells. Nat Methods 8, 821–827 (2011).

37. Ogris, C., Guala, D., Helleday, T. & Sonnhammer, E.L. A novel method for crosstalk analysis of biological networks: improving accuracy of pathway annotation. Nucleic Acids Res 45, e8 (2017).

