## Supplementary material for "Combined mRNA and protein single cell analysis in a dynamic cellular system using SPARC": Supp Figure 1-8

Figure S1

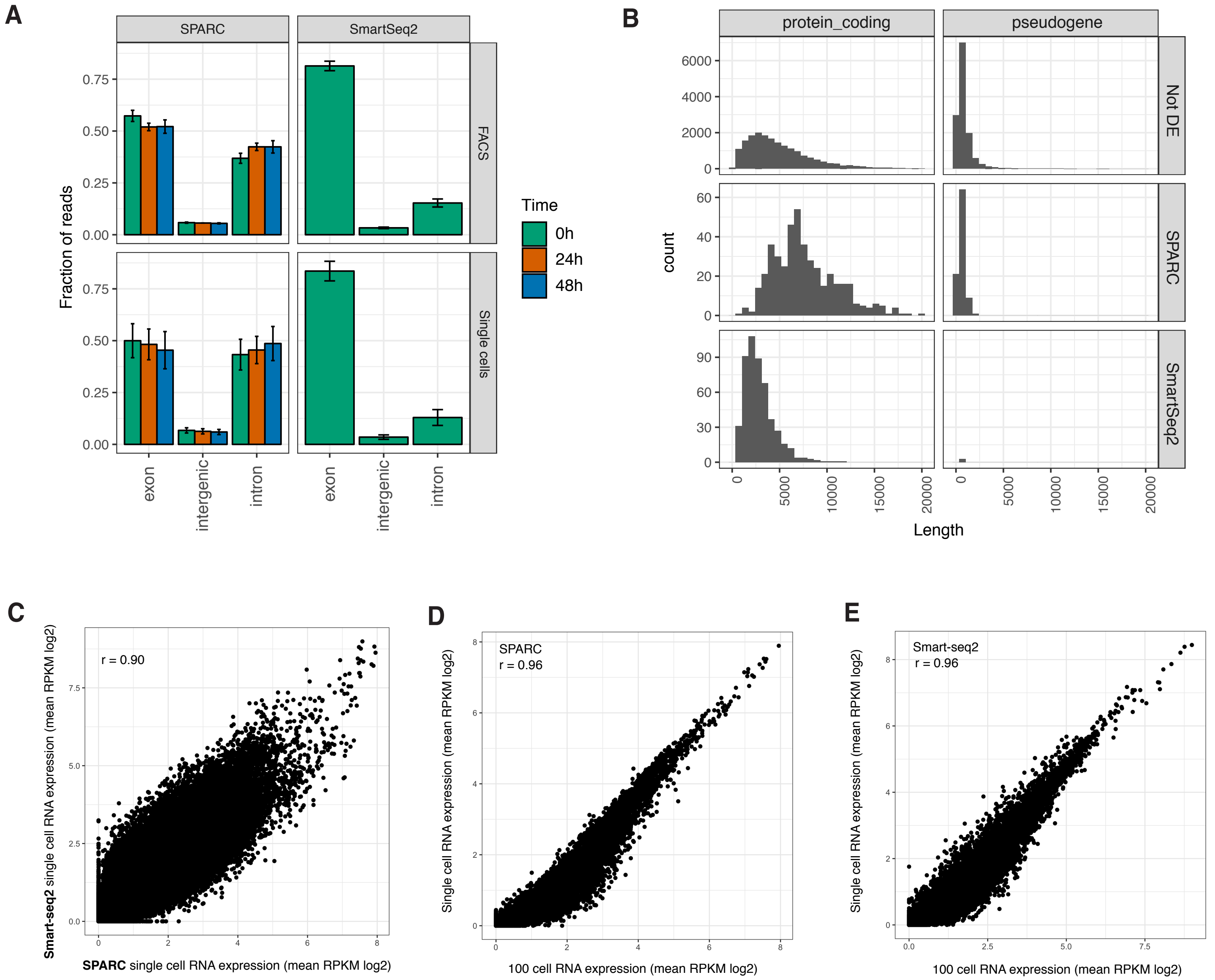
A
SPARC
SmartSeq2
FACS
Single cells
Fraction of reads
0.00
0.25
0.50
0.75
exon
intergenic
intron
Time
0h
24h
48h
B
protein_coding
pseudogene
Not DE
SPARC
SmartSeq2
count
Length
0
5000
10000
15000
20000
C
r = 0.90
Smart-seq2 single cell RNA expression (mean RPKM log2)
SPARC single cell RNA expression (mean RPKM log2)
D
SPARC
r = 0.96
Single cell RNA expression (mean RPKM log2)
100 cell RNA expression (mean RPKM log2)
E
Smart-seq2
r = 0.96
Single cell RNA expression (mean RPKM log2)
100 cell RNA expression (mean RPKM log2)

**Figure S2: mRNA (log RPKM) and protein (Cq) expression in cells measured at 0h(green), 24h (orange) and 48h (blue). Violin plots (single cells), dots (100 cell control).**

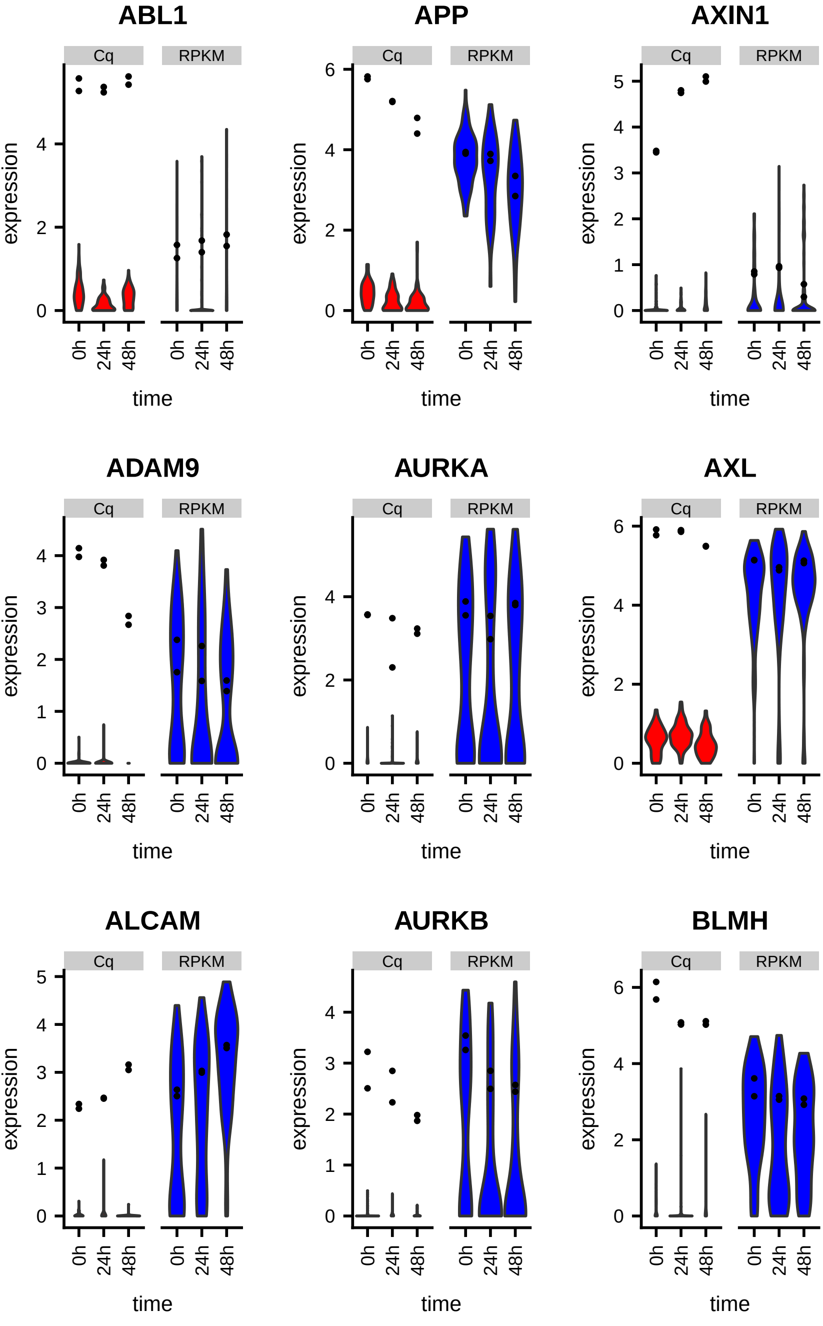

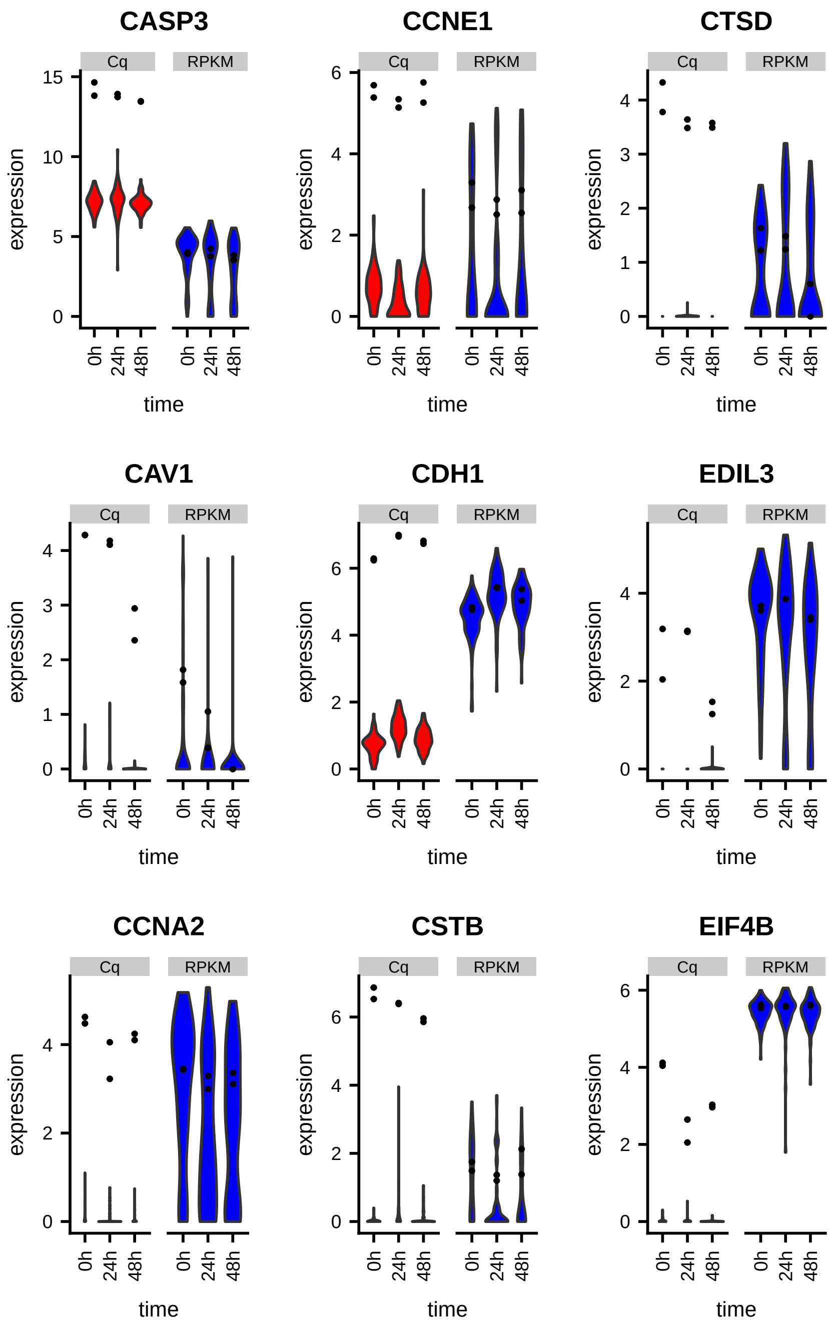
CASP3
CCNE1
CTSD
CAV1
CDH1
EDIL3
CCNA2
CSTB
EIF4B
Cq
RPKM
expression
time
0h
24h
48h

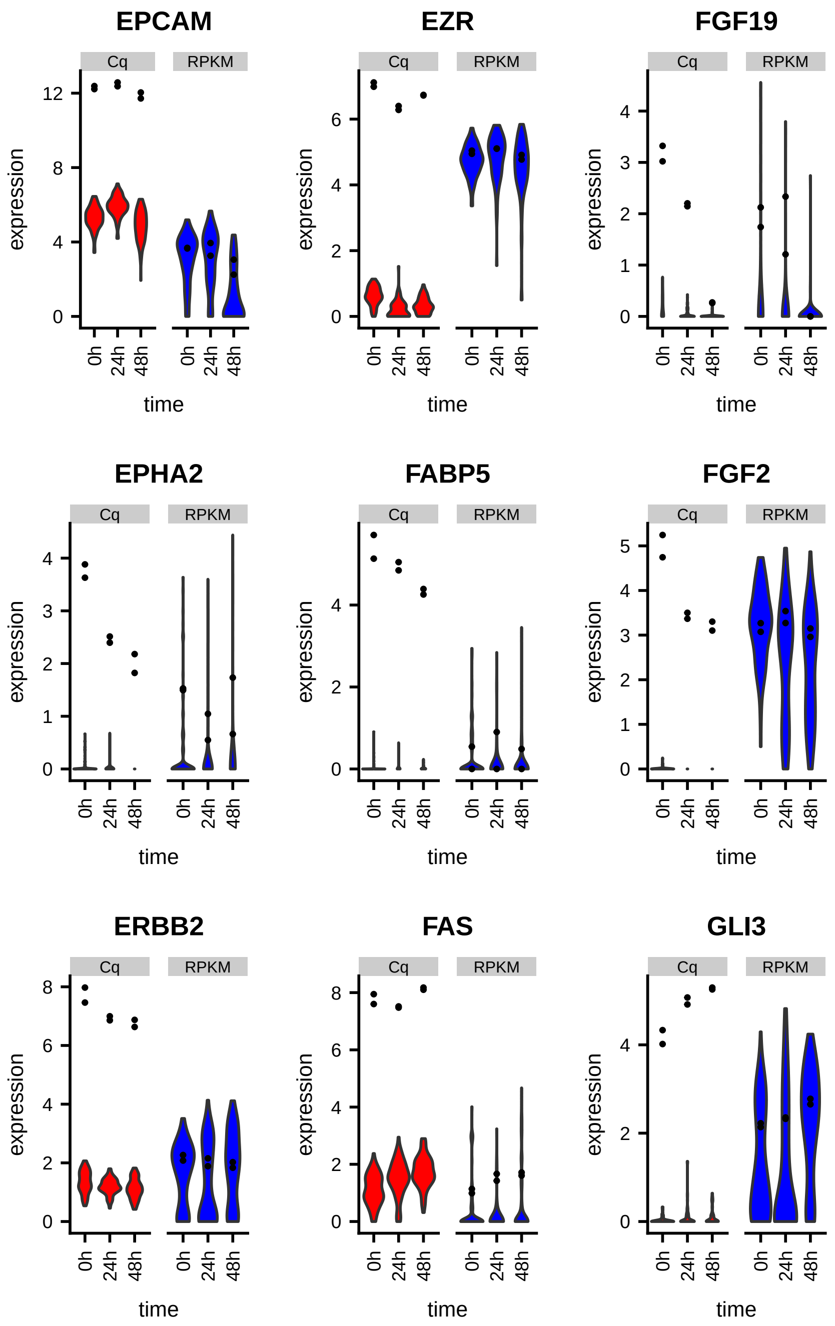

EPCAM
EZR
FGF19
EPHA2
FABP5
FGF2
ERBB2
FAS
GLI3
Cq
RPKM
expression
time
0h
24h
48h

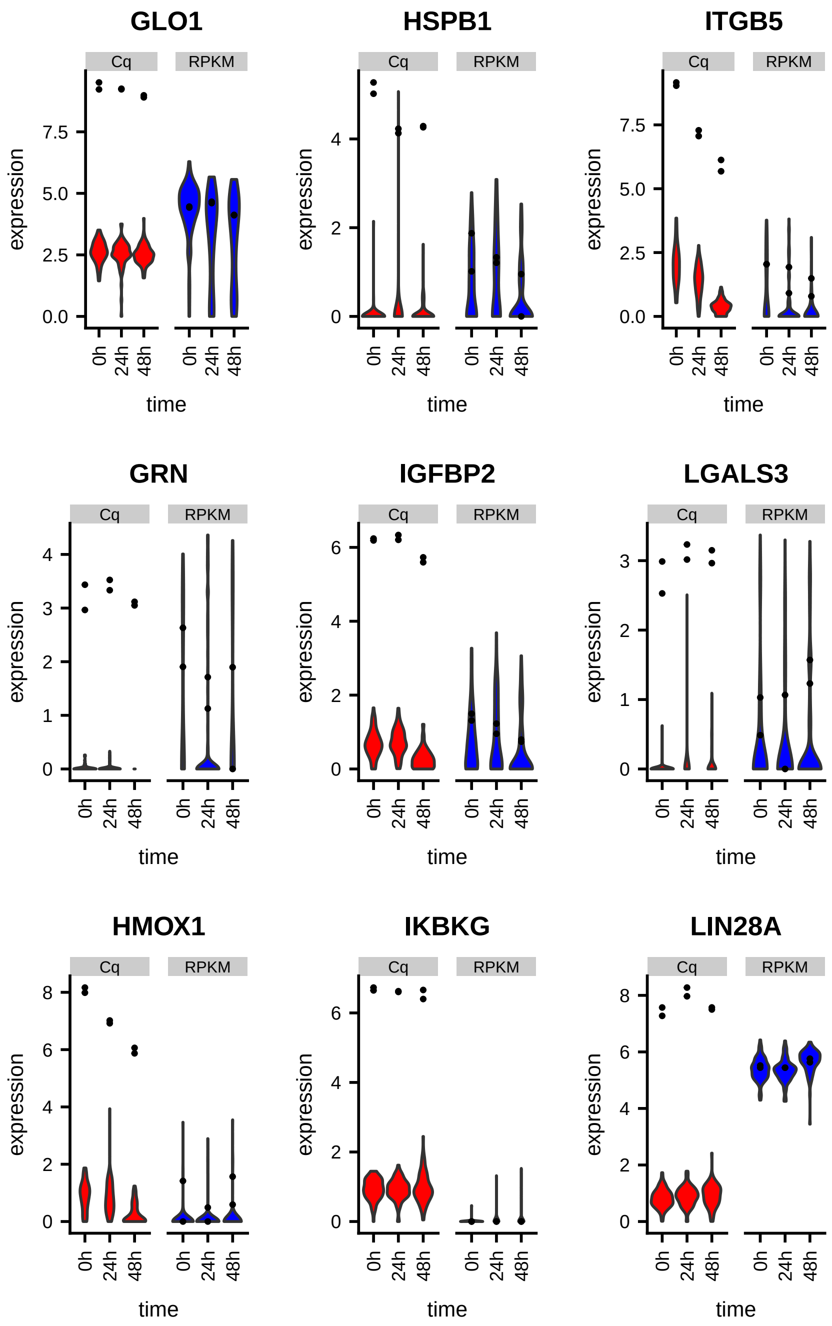
GLO1
HSPB1
ITGB5
GRN
IGFBP2
LGALS3
HMOX1
IKBKG
LIN28A
Cq
RPKM
expression
time
0h
24h
48h

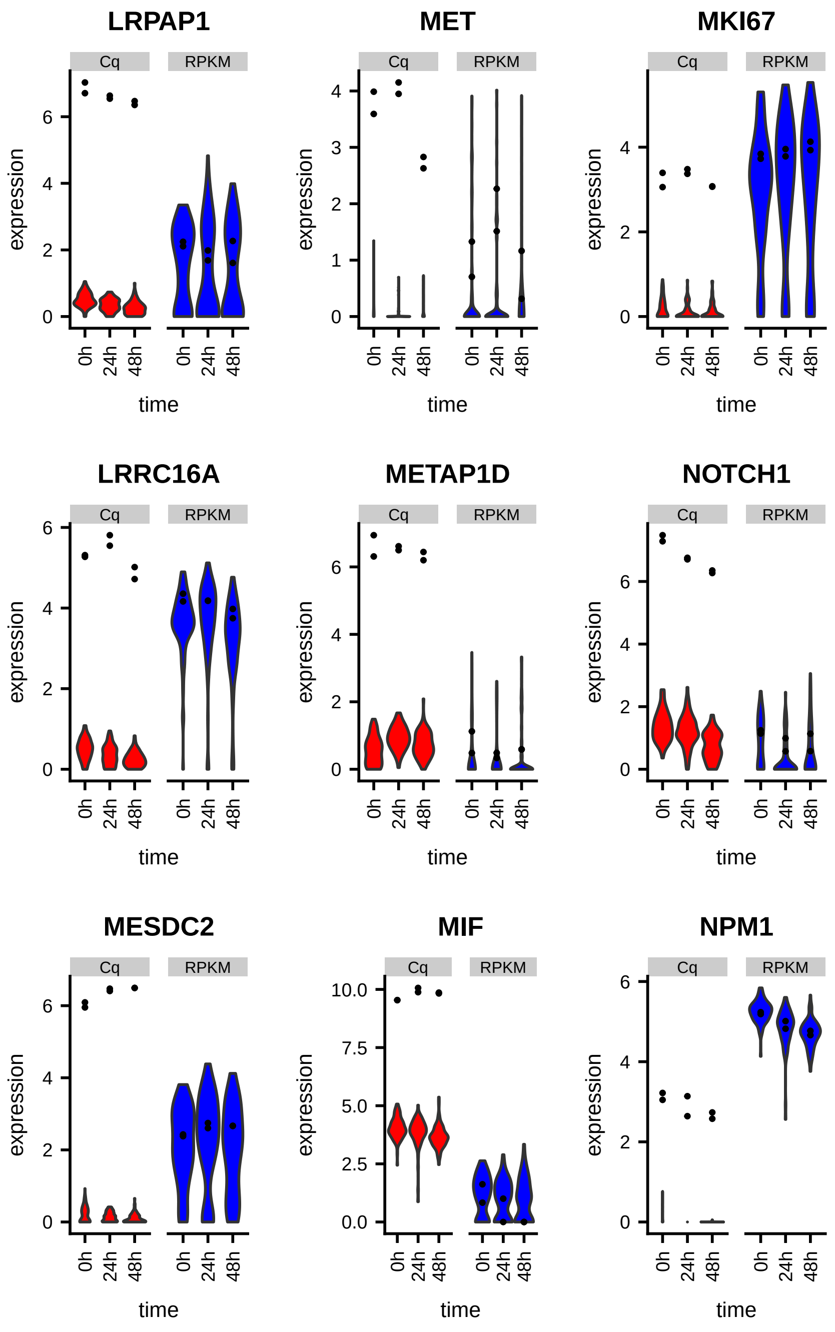

LRPAP1
MET
MKI67
LRRC16A
METAP1D
NOTCH1
MESDC2
MIF
NPM1
Cq
RPKM
expression
time
0h
24h
48h

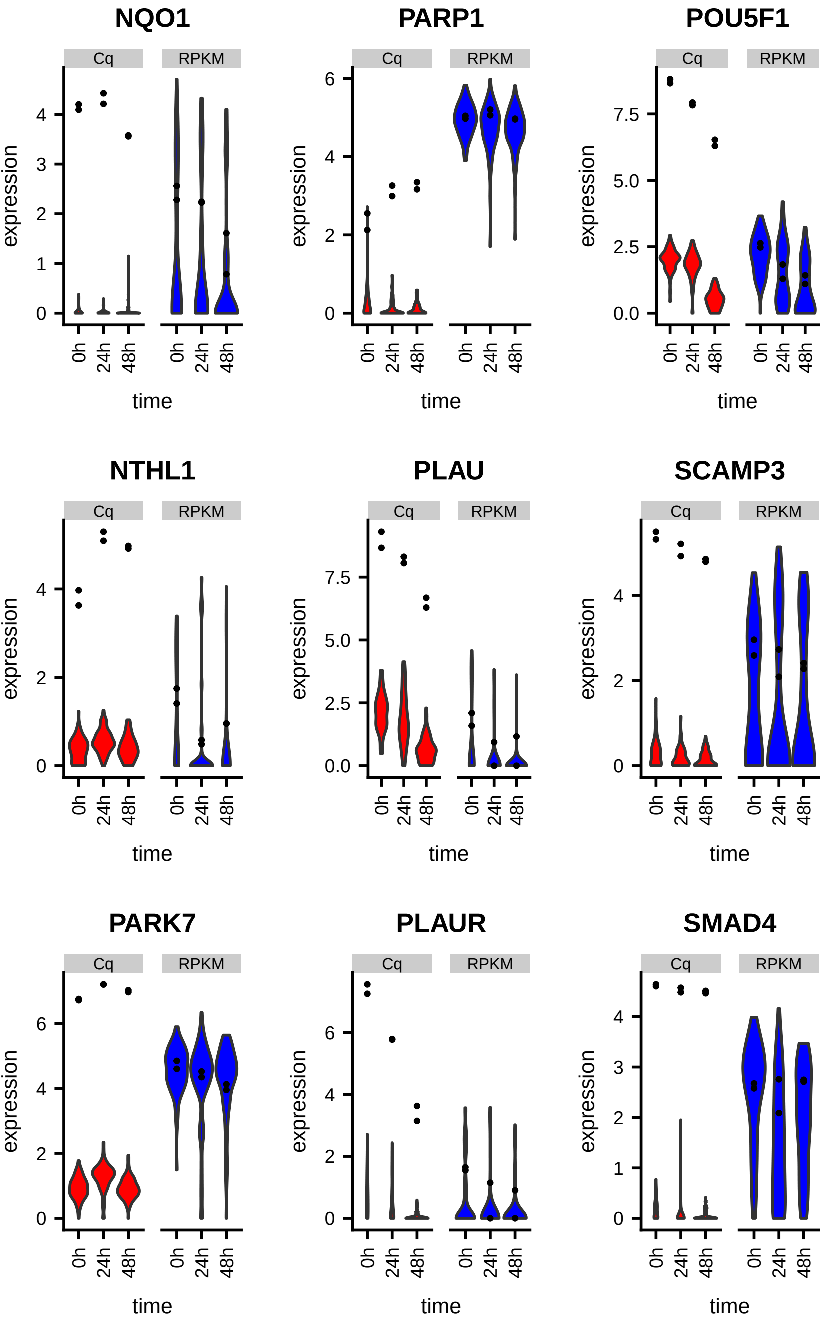
NQO1
PARP1
POU5F1
NTHL1
PLAU
SCAMP3
PARK7
PLAUR
SMAD4
Cq
RPKM
expression
time
0h
24h
48h

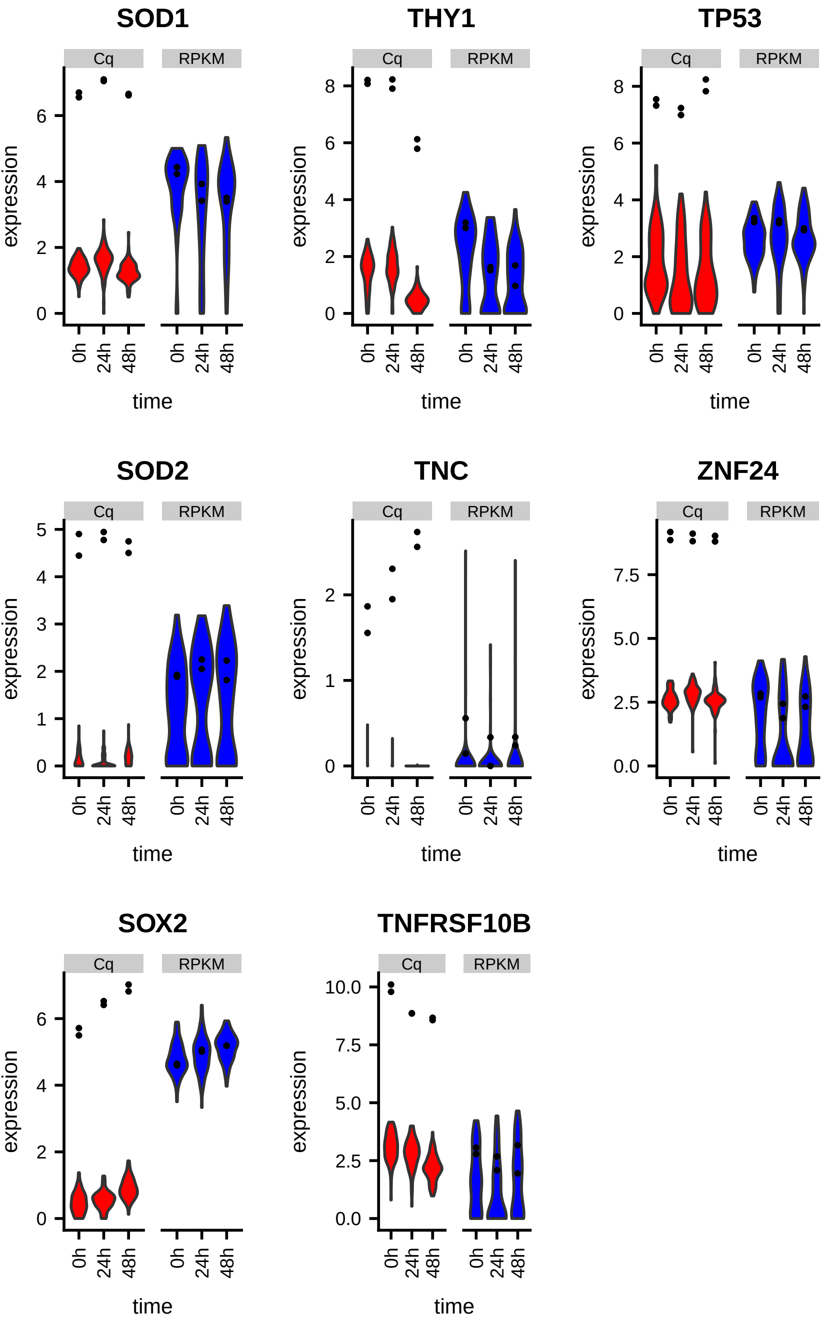
SOD1
THY1
TP53
SOD2
TNC
ZNF24
SOX2
TNFRSF10B
Cq
RPKM
expression
time
0h
24h
48h

Figure S3

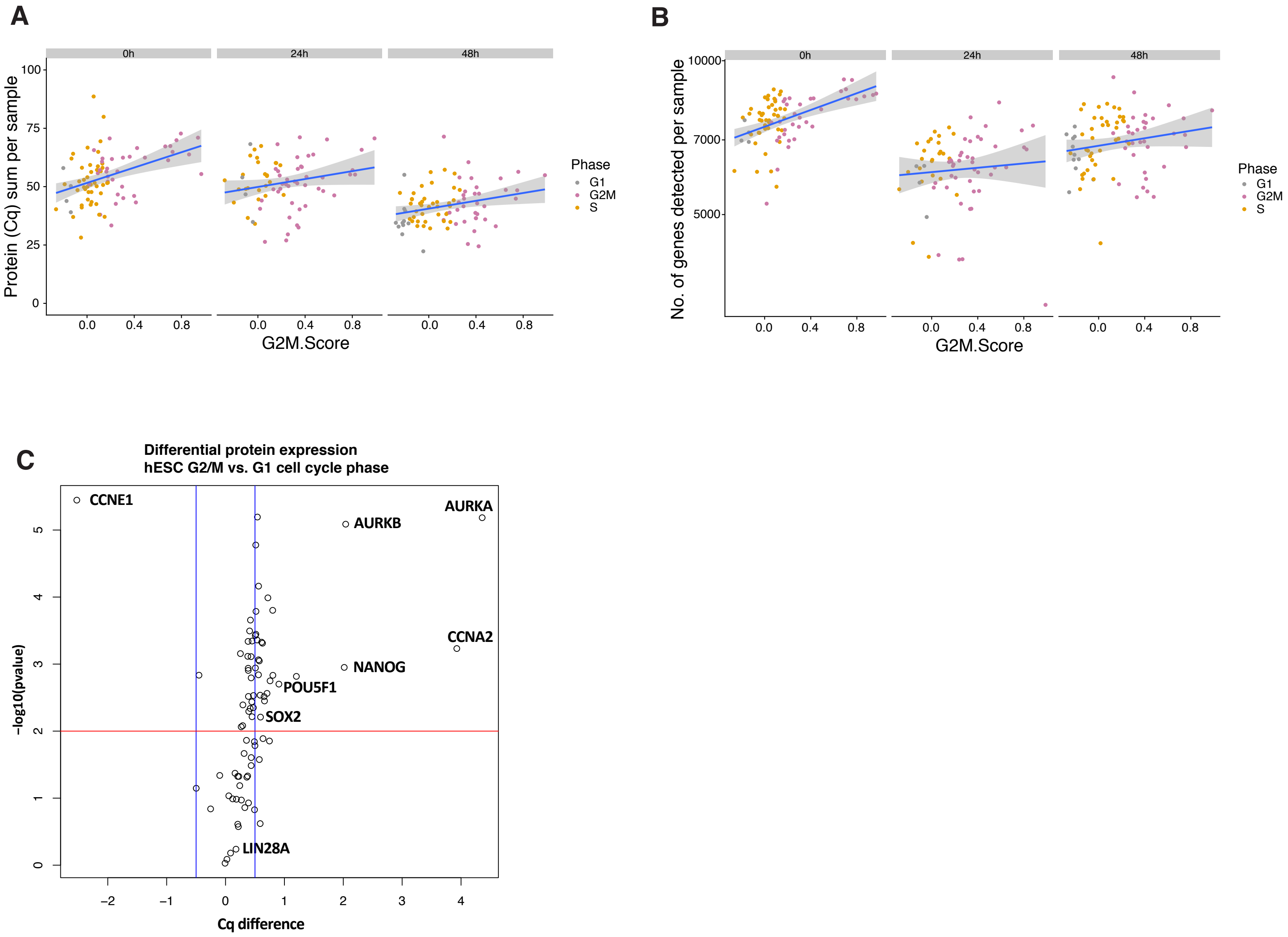
A
0h
24h
48h
Protein (Cq) sum per sample
G2M.Score
Phase
G1
G2M
S
B
0h
24h
48h
No. of genes detected per sample
G2M.Score
Phase
G1
G2M
S
C
Differential protein expression
hESC G2/M vs. G1 cell cycle phase
CCNE1
AURKB
AURKA
CCNA2
NANOG
POU5F1
SOX2
LIN28A
-log10(pvalue)
Cq difference

**Figure S4: mRNA (log RPKM) and protein (Cq) expression in cells measured at 0h (green), 24h (orange) and 48h (blue).**

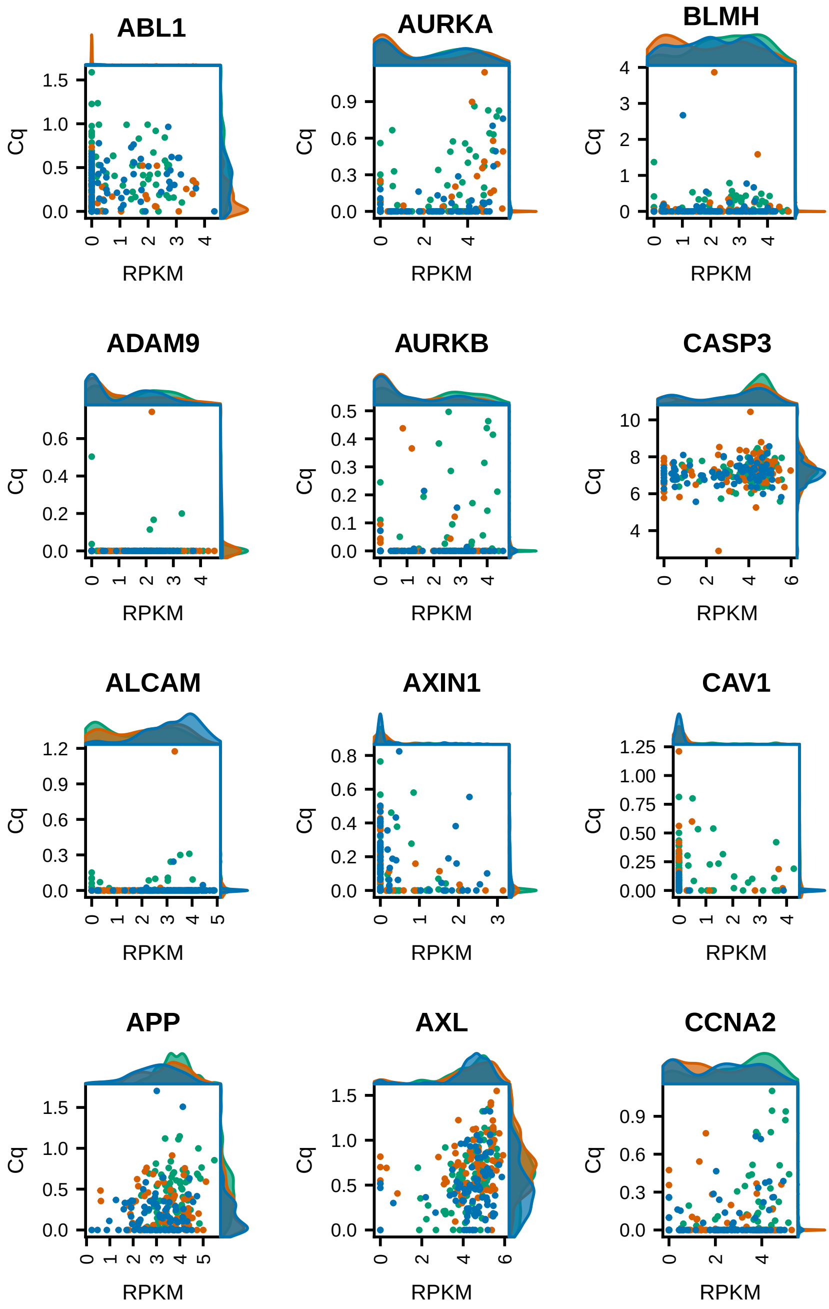

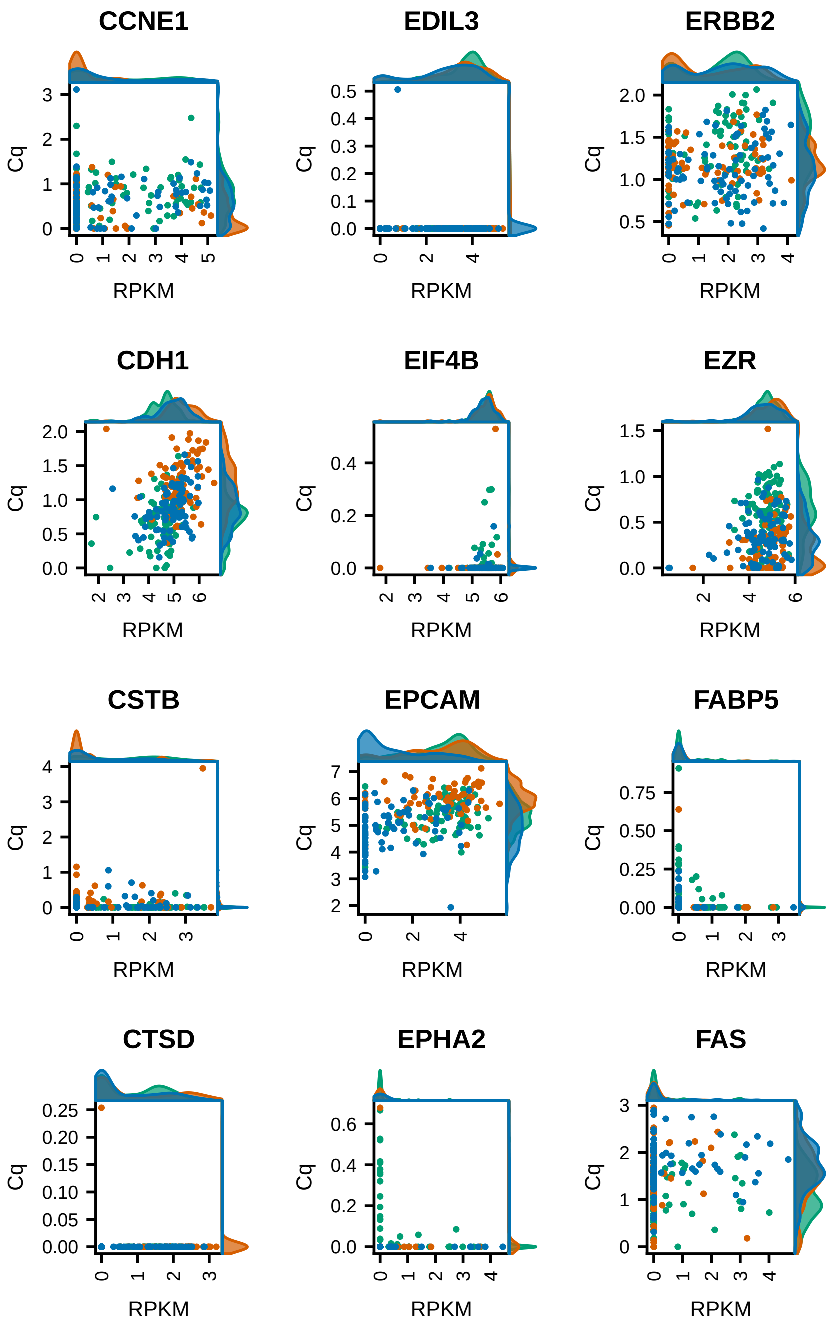

CCNE1
EDIL3
ERBB2
CDH1
EIF4B
EZR
CSTB
EPCAM
FABP5
CTSD
EPHA2
FAS
Cq
RPKM

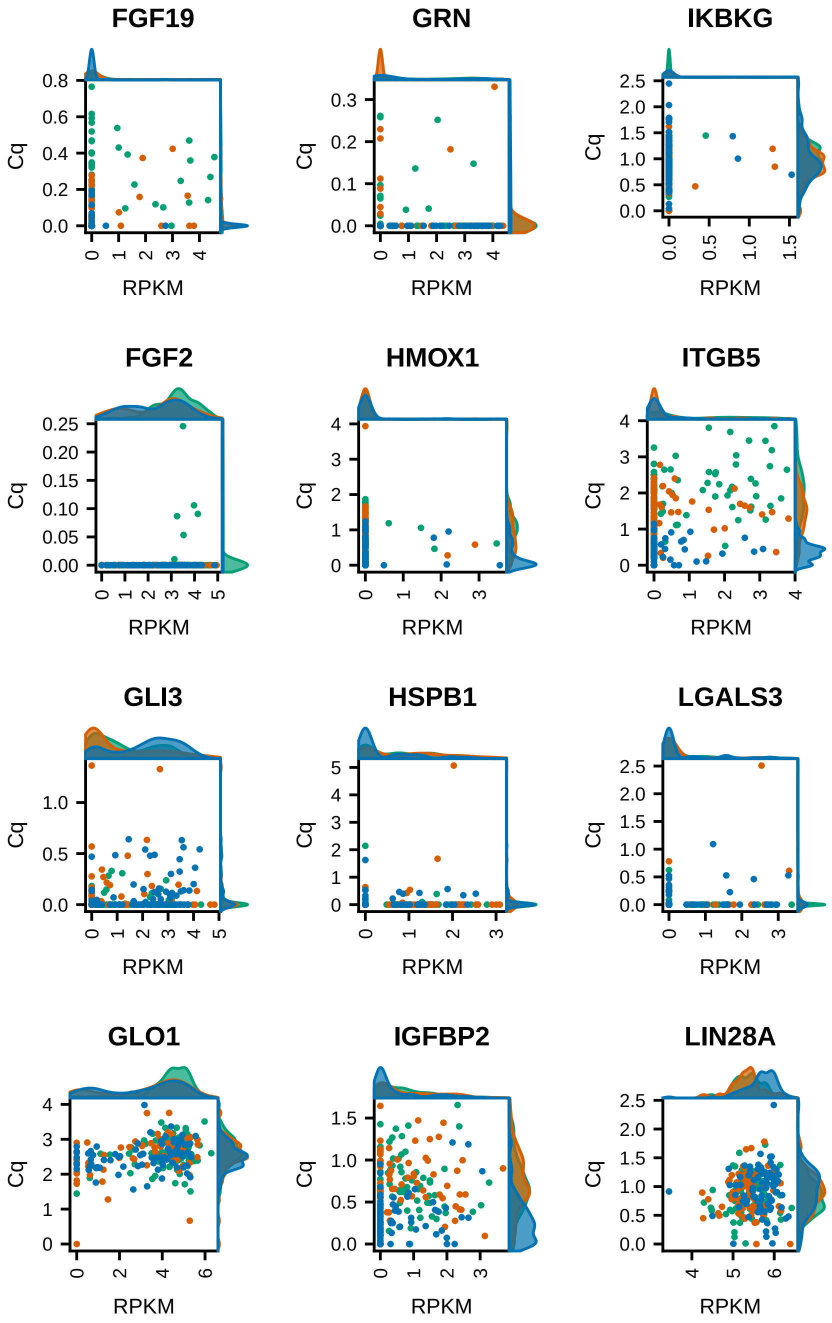
FGF19
GRN
IKBKG
FGF2
HMOX1
ITGB5
GLI3
HSPB1
LGALS3
GLO1
IGFBP2
LIN28A
Cq
RPKM

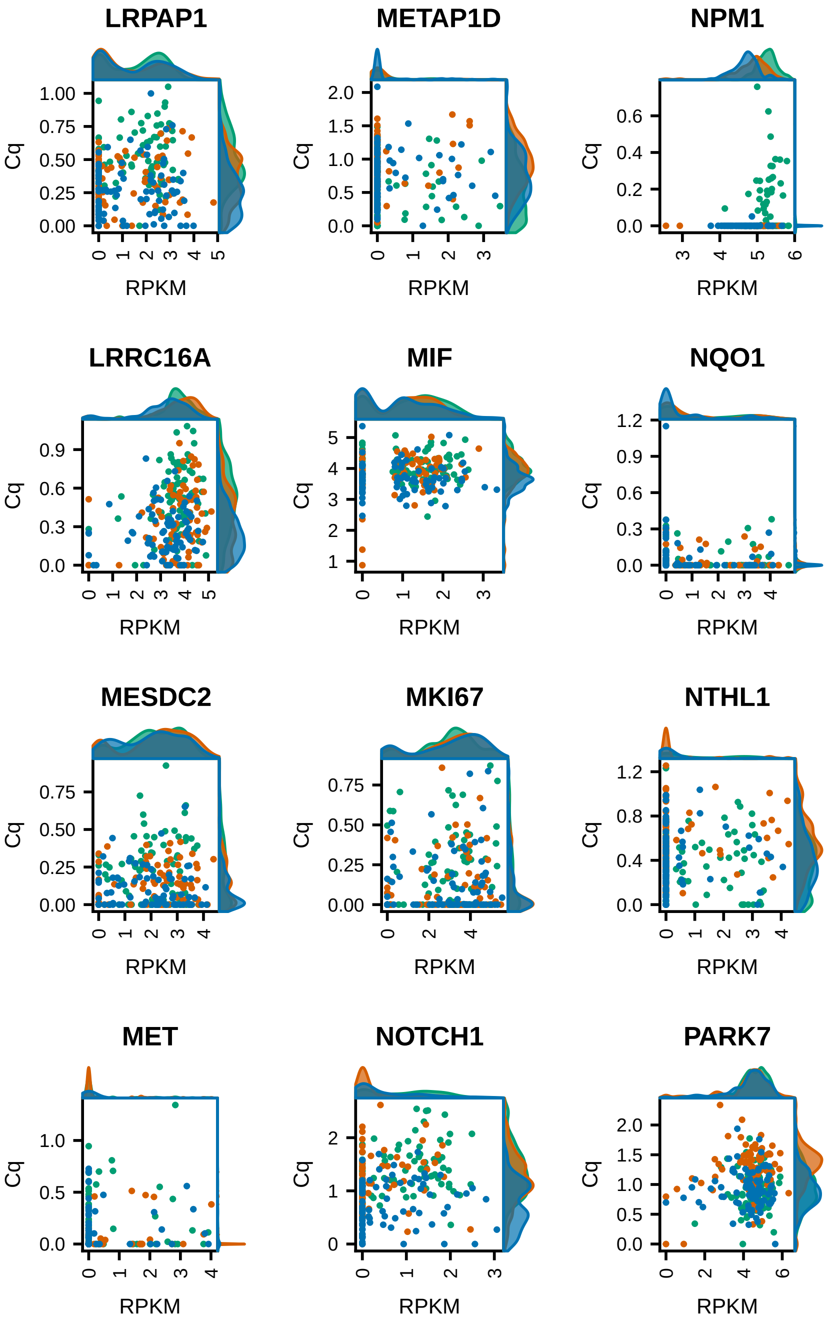
LRPAP1
METAP1D
NPM1
LRRC16A
MIF
NQO1
MESDC2
MKI67
NTHL1
MET
NOTCH1
PARK7
Cq
RPKM

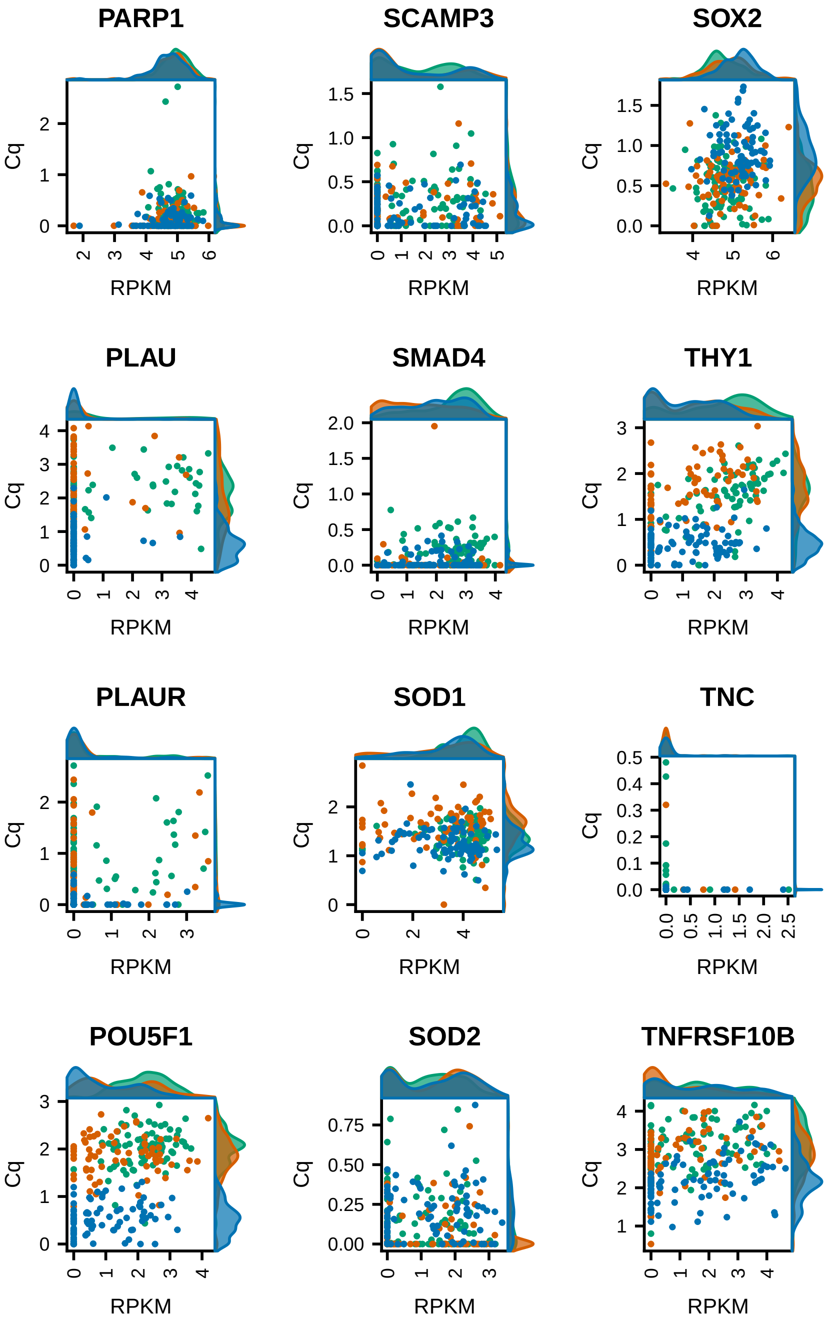
PARP1
SCAMP3
SOX2
PLAU
SMAD4
THY1
PLAUR
SOD1
TNC
POU5F1
SOD2
TNFRSF10B
Cq
RPKM

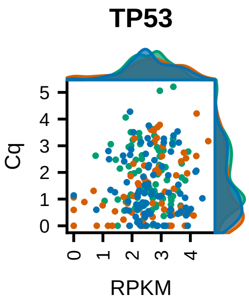
TP53
Cq
5
4
3
2
1
0
0
1
2
3
4
RPKM

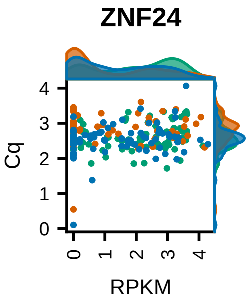
ZNF24
Cq
4
3
2
1
0
0
1
2
3
4
RPKM

**Figure S5: mRNA and protein expression in cells ordered by pseudotime$_{RNA}$**

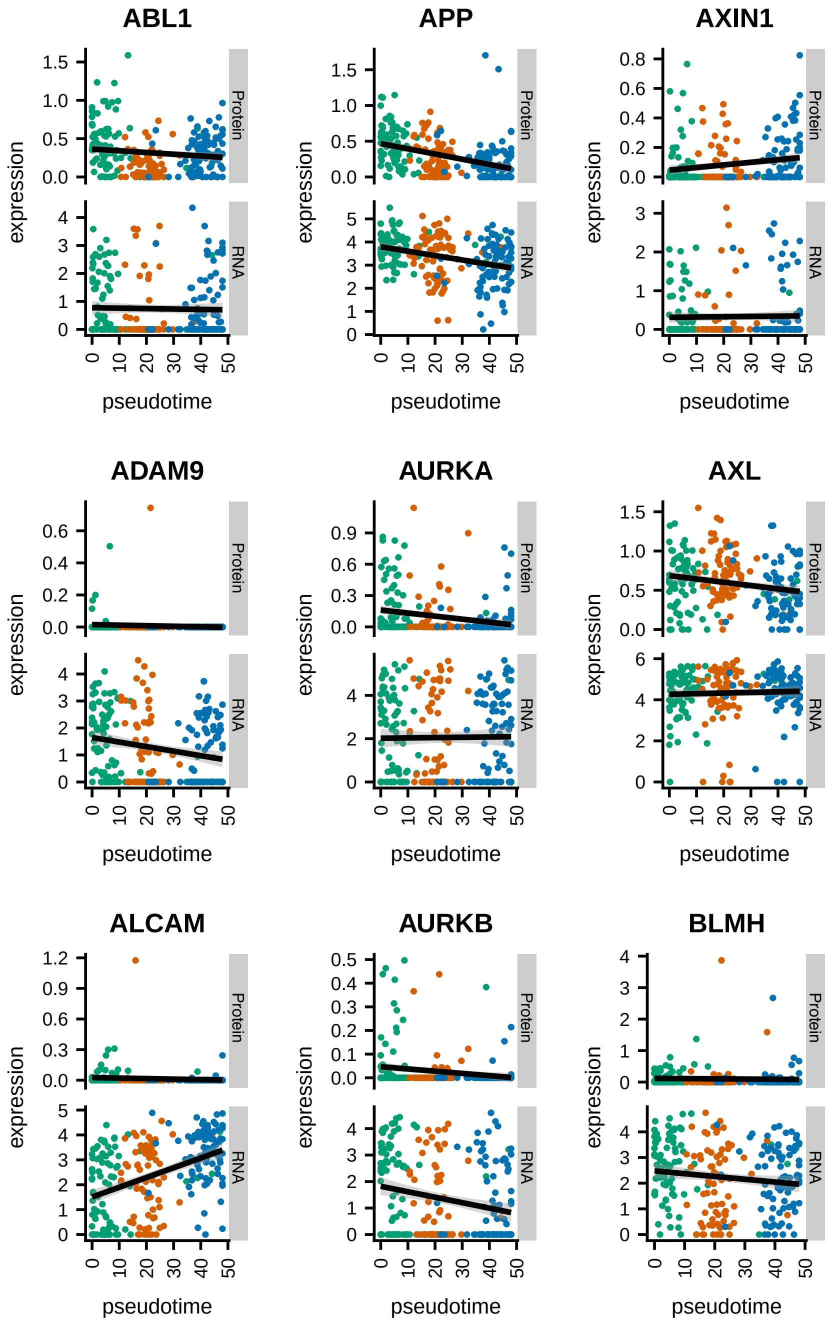

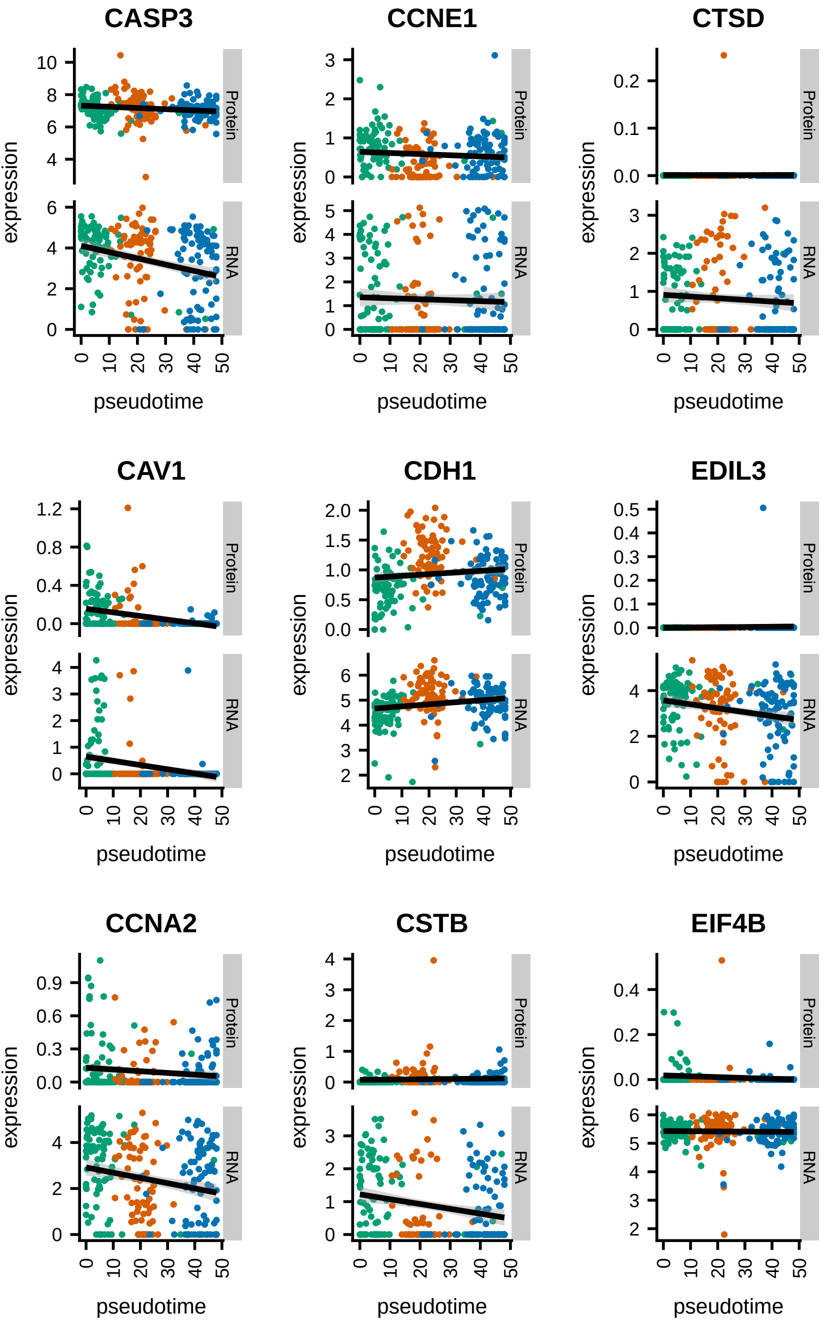

CASP3
CCNE1
CTSD
CAV1
CDH1
EDIL3
CCNA2
CSTB
EIF4B
Protein
RNA
expression
pseudotime

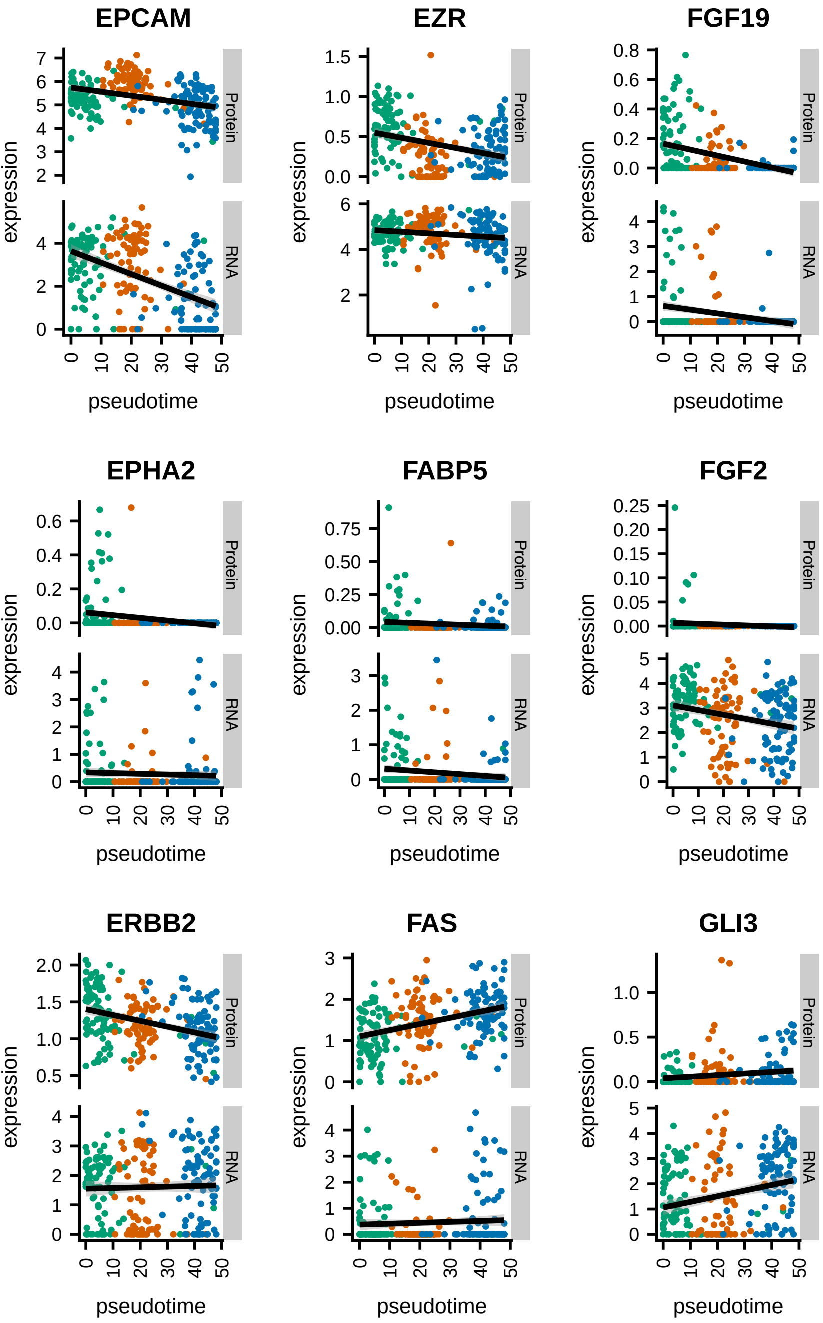

EPCAM
EZR
FGF19
EPHA2
FABP5
FGF2
ERBB2
FAS
GLI3
expression
pseudotime
Protein
RNA

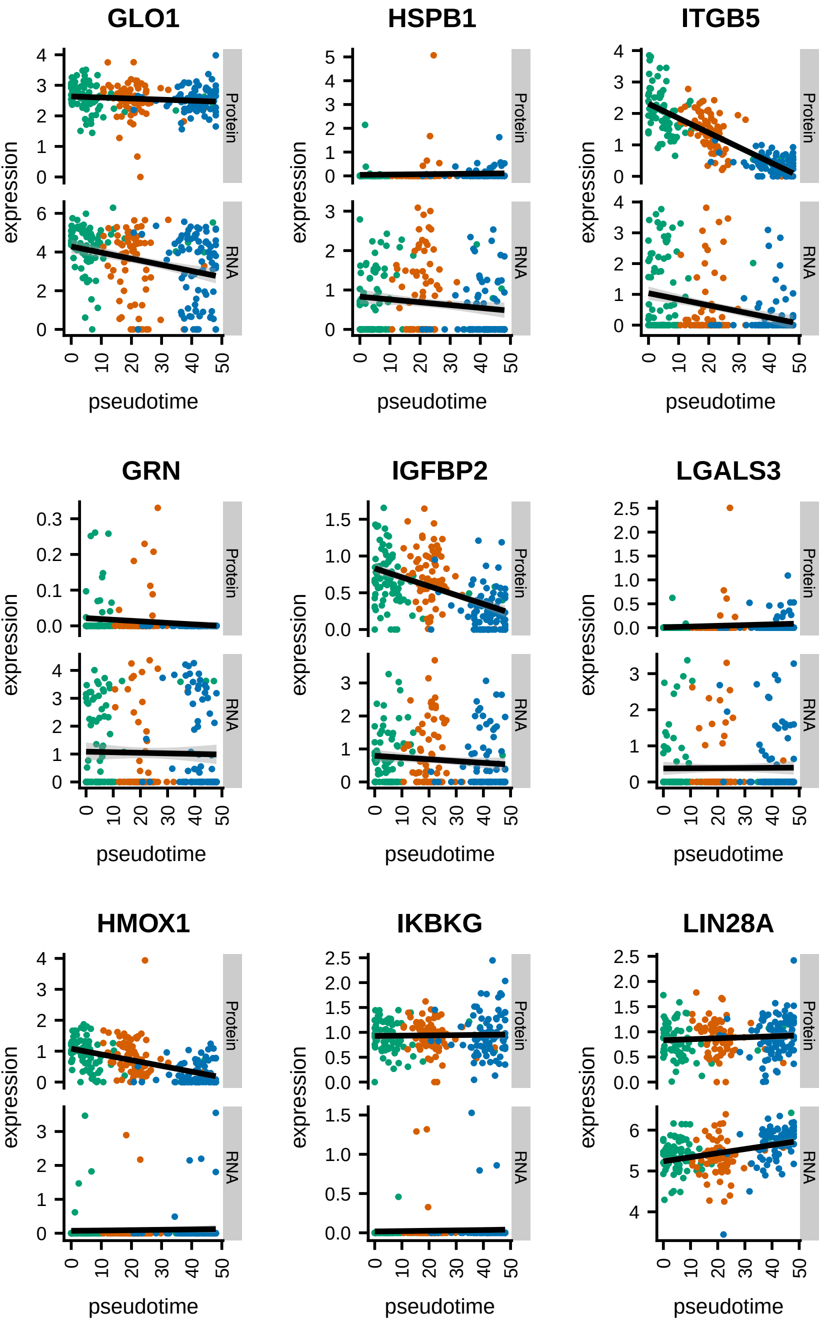
GLO1
HSPB1
ITGB5
GRN
IGFBP2
LGALS3
HMOX1
IKBKG
LIN28A
expression
pseudotime
Protein
RNA

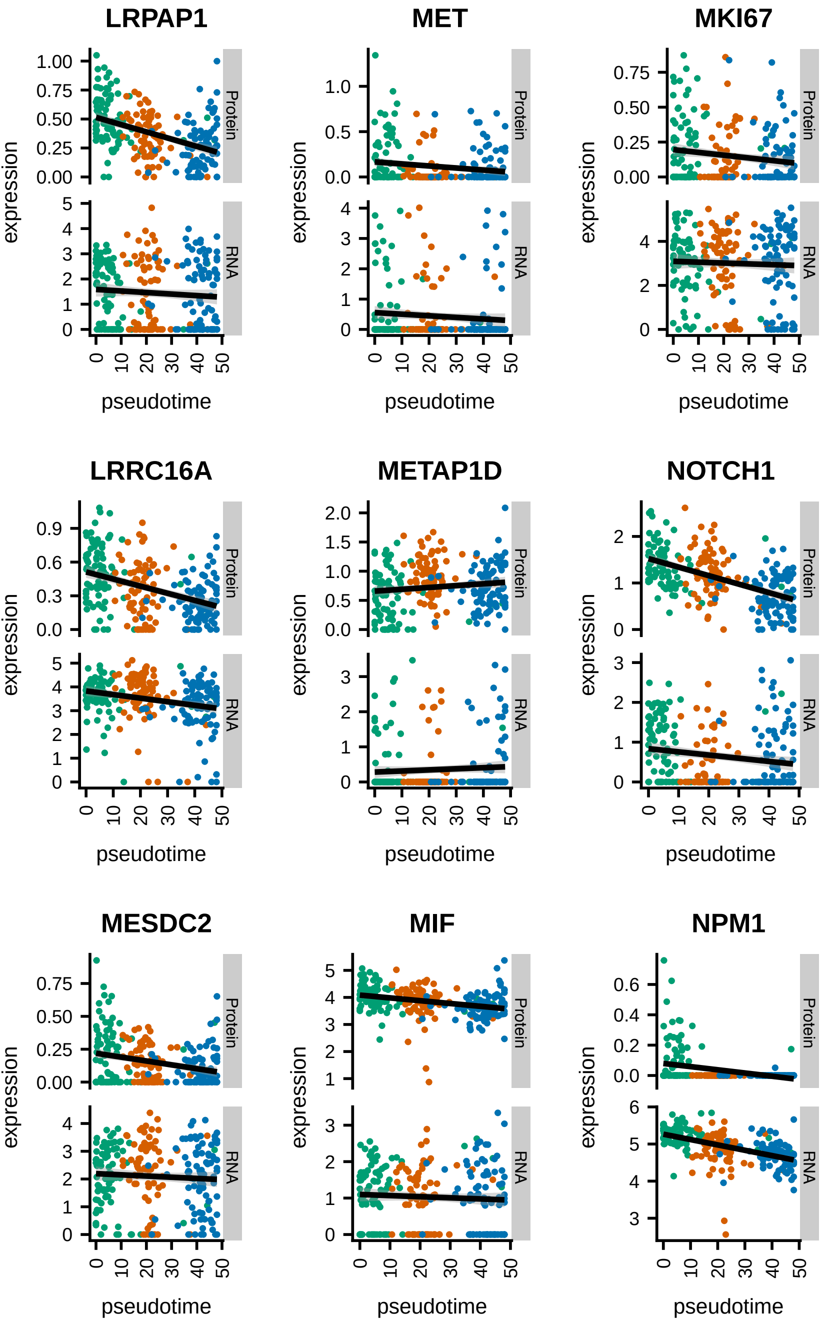

LRPAP1
MET
MKI67
LRRC16A
METAP1D
NOTCH1
MESDC2
MIF
NPM1
Protein
RNA
expression
pseudotime

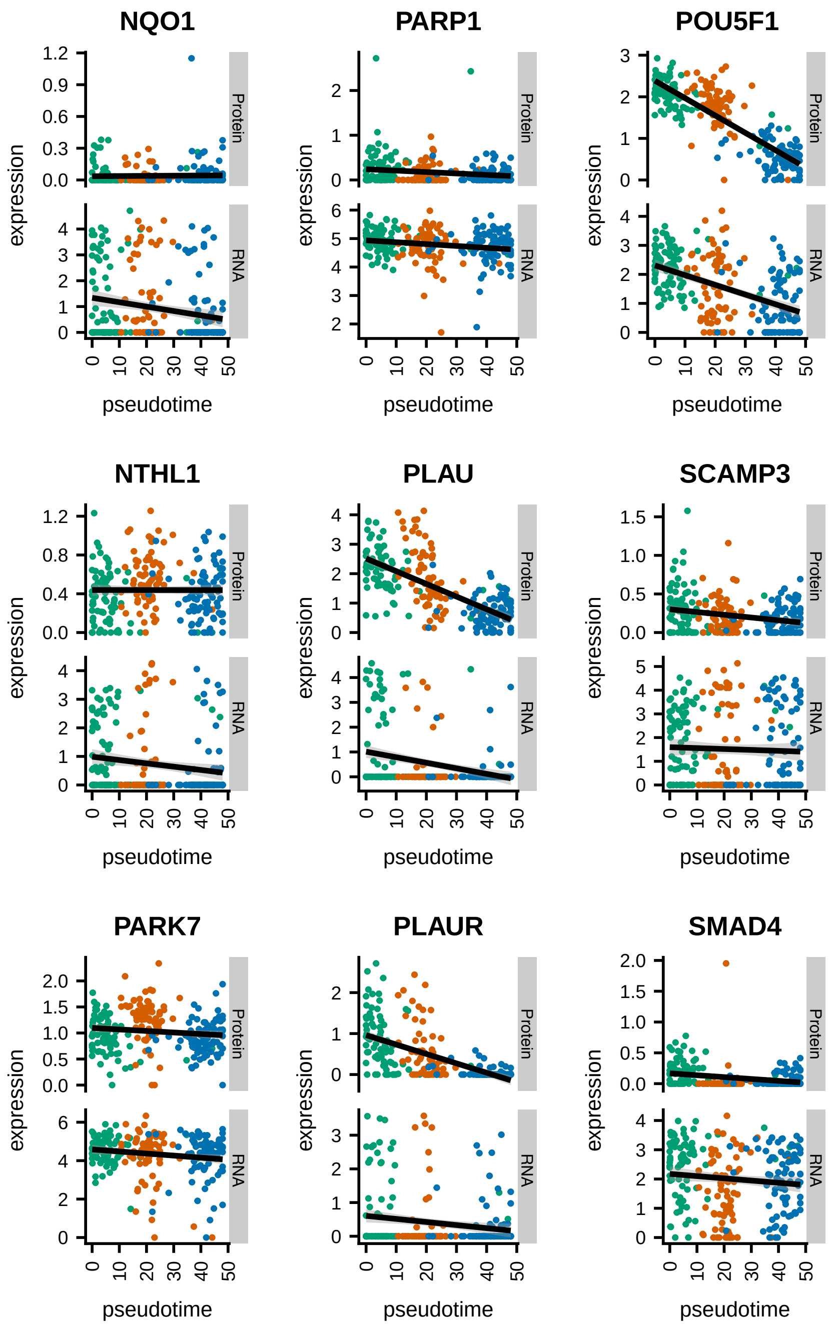
NQO1
PARP1
POU5F1
NTHL1
PLAU
SCAMP3
PARK7
PLAUR
SMAD4
expression
pseudotime
Protein
RNA

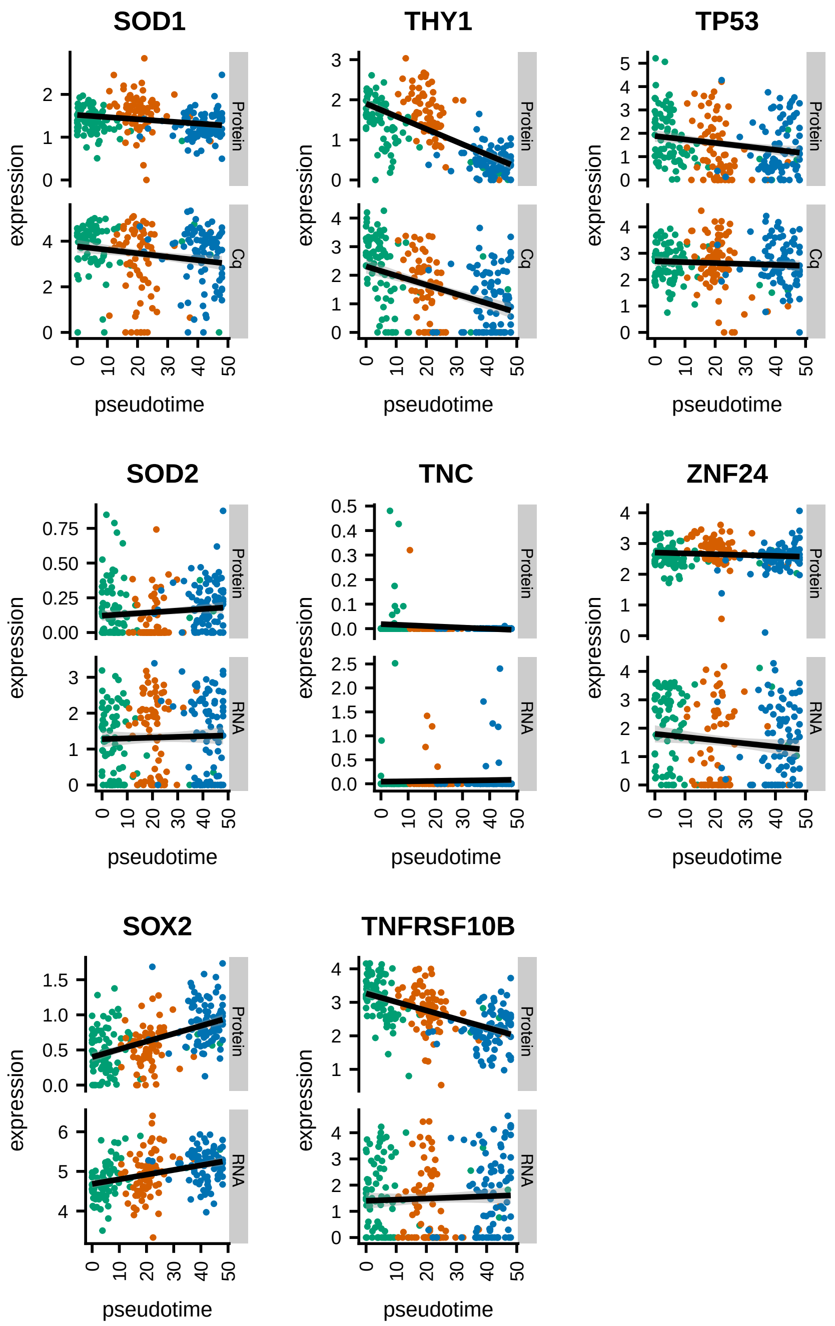

SOD1
THY1
TP53
SOD2
TNC
ZNF24
SOX2
TNFRSF10B
expression
pseudotime
Protein
Cq
RNA

Figure S6

A

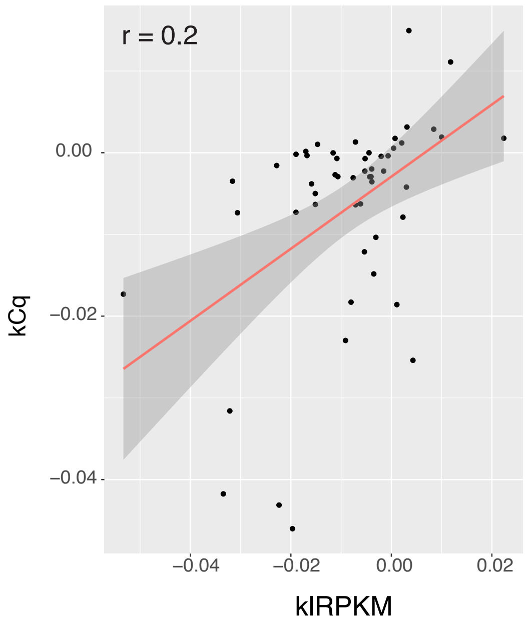

Figure S7

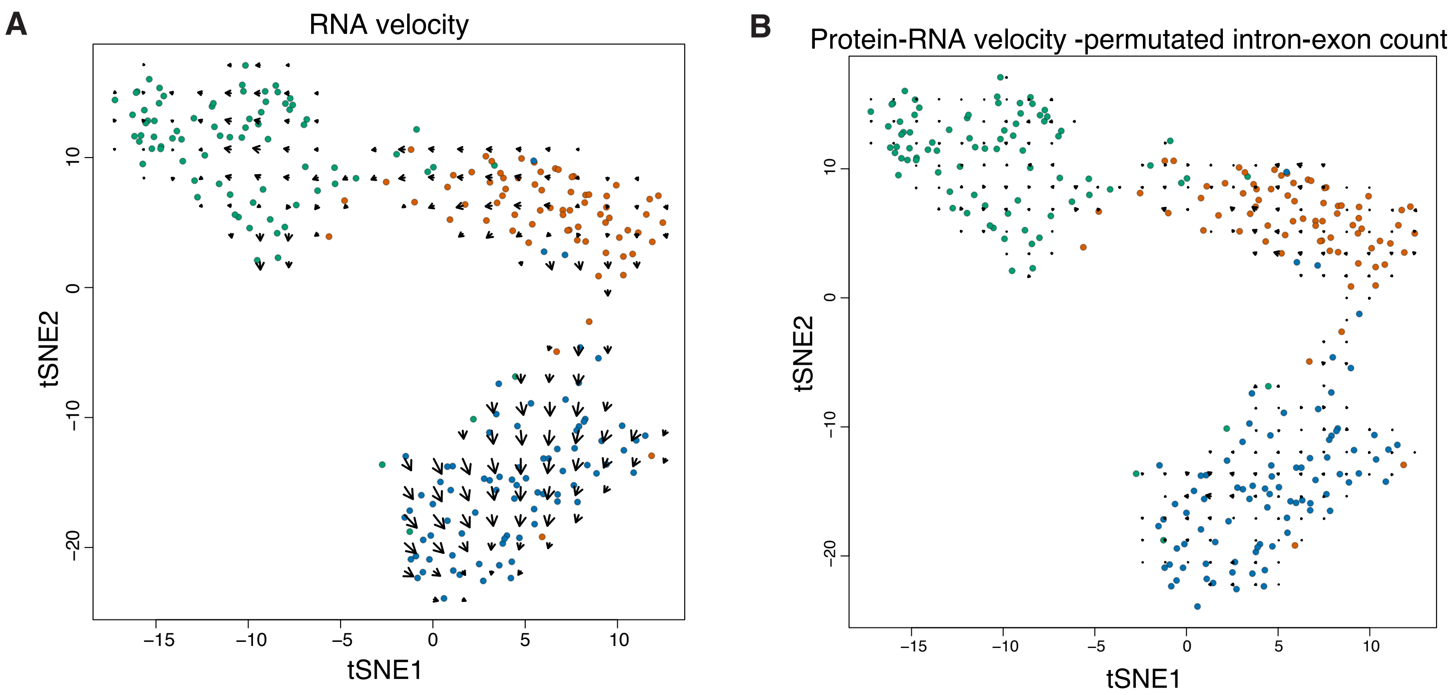
A
RNA velocity
tSNE2
tSNE1
10
0
−10
−20
−15
−10
−5
0
5
10
B
Protein-RNA velocity -permutated intron-exon count
tSNE2
tSNE1
10
0
−10
−20
−15
−10
−5
0
5
10

### Figure S8

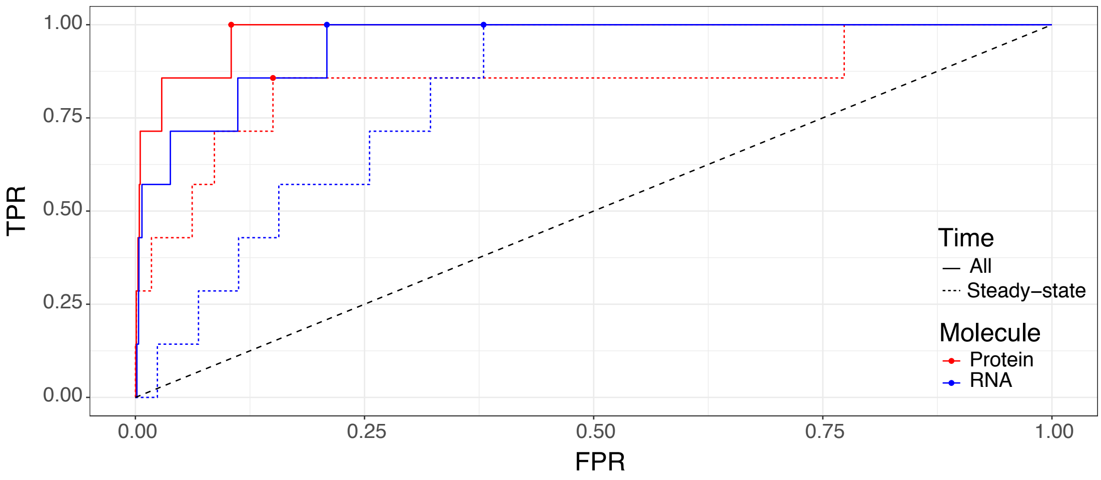

TPR
1.00
0.75
0.50
0.25
0.00
0.00
0.25
0.50
0.75
1.00
FPR
Time
All
Steady–state
Molecule
Protein
RNA
